# SARS-CoV-2 and SARS-CoV spike-mediated cell-cell fusion differ in the requirements for receptor expression and proteolytic activation

**DOI:** 10.1101/2020.07.25.221135

**Authors:** Bojan F. Hörnich, Anna K. Großkopf, Sarah Schlagowski, Matthias Tenbusch, Hannah Kleine-Weber, Frank Neipel, Christiane Stahl-Hennig, Alexander S. Hahn

## Abstract

The severe acute respiratory syndrome-related coronavirus 2 (SARS-CoV-2) infects cells through interaction of its spike protein (SARS2-S) with Angiotensin-converting enzyme 2 (ACE2) and activation by proteases, in particular transmembrane protease serine 2 (TMPRSS2). Viruses can also spread through fusion of infected with uninfected cells. We compared the requirements of ACE2 expression, proteolytic activation, and the sensitivity to inhibitors for SARS2-S-mediated and SARS-CoV-S(SARS1-S)-mediated cell-cell fusion. SARS2-S-driven fusion was moderately increased by TMPRSS2 and strongly by ACE2, while SARS1-S-driven fusion was strongly increased by TMPRSS2 and less so by ACE2 expression. In contrast to SARS1-S, SARS2-S-mediated cell-cell fusion was efficiently activated by Batimastat-sensitive metalloproteases. Mutation of the S1/S2 proteolytic cleavage site reduced effector-target-cell fusion when ACE2 or TMPRSS2 were limiting and rendered SARS2-S-driven cell-cell fusion more dependent on TMPRSS2. When both ACE2 and TMPRSS2 were abundant, initial target-effector-cell fusion was unaltered compared to wt SARS2-S, but syncytia remained smaller. Mutation of the S2’ site specifically abrogated activation by TMPRSS2 for both cell-cell fusion and SARS2-S-driven pseudoparticle entry but still allowed for activation by metalloproteases for cell-cell fusion and by cathepsins for particle entry. Finally, we found that the TMPRSS2 inhibitor Bromhexine was unable to reduce TMPRSS2-activated cell-cell fusion by SARS1-S and SARS2-S as opposed to the inhibitor Camostat. Paradoxically, Bromhexine enhanced cell-cell fusion in the presence of TMPRSS2, while its metabolite Ambroxol exhibited inhibitory activity in some conditions. On Calu-3 lung cells, Ambroxol weakly inhibited SARS2-S-driven lentiviral pseudoparticle entry, and both substances exhibited a dose-dependent trend towards weak inhibition of authentic SARS-CoV-2.

**IMPORTANCE:** Cell-cell fusion allows the virus to infect neighboring cells without the need to produce free virus and contributes to tissue damage by creating virus-infected syncytia. Our results demonstrate that the S2’ cleavage site is essential for activation by TMPRSS2 and unravel important differences between SARS-CoV and SARS-CoV-2, among those greater dependence of SARS-CoV-2 on ACE2 expression and activation by metalloproteases for cell-cell fusion. Bromhexine, reportedly an inhibitor of TMPRSS2, is currently tested in clinical trials against coronavirus disease 2019. Our results indicate that Bromhexine enhances fusion in some conditions. We therefore caution against use of Bromhexine in higher dosage until its effects on SARS-CoV-2 spike activation are better understood. The related compound Ambroxol, which similarly to Bromhexine is clinically used as an expectorant, did not exhibit activating effects on cell-cell fusion. Both compounds exhibited weak inhibitory activity against SARS-CoV-2 infection at high concentrations, which might be clinically attainable for Ambroxol.

## INTRODUCTION

The coronavirus disease 2019 (COVID-19) disease spectrum is caused by the severe acute respiratory syndrome-related coronavirus 2 (SARS-CoV-2), which was first identified in patients with pneumonia of unknown origin in the city of Wuhan, China (1). While first characterized as a pneumonia, COVID-19 probably affects a number of organ systems (2–4). SARS-CoV-2 was shown to use the Angiotensin-converting enzyme 2 (ACE2) receptor, which was previously described as receptor for the closely related severe acute respiratory syndrome-related coronavirus (SARS-CoV) (5), for the infection of human cells (1, 6, 7). For the proteolytic activation of the viral spike protein, a prerequisite for fusion activity of coronaviruses (reviewed in (8)), the transmembrane protease serine 2 (TMPRSS2) (7, 9) as well as the related TMPRSS4 (2) were reported to be of critical importance. In addition, TMPRSS2 was demonstrated to colocalize with the ACE2 receptor (10) and therefore may be biologically particularly relevant.

Depending on the cell type, SARS-CoV-2 spike(SARS2-S)-driven entry can also occur through endocytotic pathways where virus-cell fusion is most likely activated by cathepsins (7). Another study reported that several members of the TMPRSS family can activate SARS2-S-mediated membrane fusion (11). The proposed mechanisms for spike priming and initiation of fusion therefore require further clarification, e.g. whether serine protease activity is required under all circumstances, or whether fusion can also occur without the action of serine proteases, where these proteases act on the spike, and whether there are differences between cell-cell and cell-particle fusion.

It was recently discovered that the polybasic S1/S2 cleavage site of SARS2-S is required for efficient infection of lung-derived cells and promotes the formation of syncytia (12). Understanding syncytium formation may be important as large syncytial elements are reported to constitute a hallmark of COVID-19-associated pathology (13). Nevertheless, the exact contribution of the two known proteolytic priming sites to cell-cell fusion and their protease usage are not entirely clear. To address these questions, we mutated the S1/S2 site as well as the S2’ site, we assessed the effects of proteolytic activation by using inhibitors of TMPRSS2 and other proteases, and we analyzed the effects of different levels of protease and receptor expression on SARS-CoV spike (SARS1-S) and SARS2-S fusion activity.

TMPRSS2, which is expressed in airway cells (14), may be amenable to specific inhibition by Bromhexine (15), a molecule normally used as an expectorant that thins phlegm and eases coughing and is widely known as a popular over-the-counter medication, which would make its repurposing for COVID-19 particularly attractive. For these or additional reasons, Bromhexine is now being tested in at least three clinical trials for efficacy against COVID-19 (NCT04355026, NCT04273763, NCT04340349). We therefore tested the effect of the TMPRSS2 inhibitor Bromhexine on spike-mediated cell-cell fusion and SARS2-S driven cell entry and compared its potency to the serine protease inhibitor Camostat. We also included Ambroxol, an active metabolite of Bromhexine in our studies (16). Ambroxol has often replaced Bromhexine as an over-the-counter medication, and the structural similarity to Bromhexine may hint at potential inhibitory effects towards TMPRSS2. Ambroxol may also exhibit weak but broad anti-viral activity as it was shown to reduce the occurrence of respiratory infections (17) and to inhibit proteolytic activation of influenza virus by triggering release of antiviral factors (18), and it is used to treat acute respiratory distress syndrome in adults and antenatally in infants (19, 20). Further, two recent preprints, one describing modulation of the ACE2-SARS2-S interaction by both Bromhexine and Ambroxol (21) and the other reporting weak inhibitory activity of Ambroxol against SARS-CoV-2 replication (22) in Vero E6 cells, point at a potential utility of these molecules in the therapy of COVID-19.

## RESULTS

### SARS2-S mediates robust fusion of 293T cells transfected with ACE2 with and without co-expression of TMPRSS2

In order to investigate the fusion mechanism of SARS-CoV-2 we generated several SARS2-S mutants (Fig. 1 A, schematic drawn after (23)). It was reported that the furin-recognition motif at the S1/S2 cleavage site of SARS2-S, which is not found in SARS1-S, plays a role in the infection of airway cells like Calu-3 but is dispensable in other cell types (11, 12). Thus, we generated a mutant, SARS2-S1/S2-mut, where the furin-recognition motif and the cleavage site were replaced by alanines (Fig 1 A). In contrast to already published S1/S2 mutants (24, 25) we did not delete the site as we suspected that this may influence protein conformation and flexibility but we mutated the proposed furin cleavage site (26) to fully abrogate processing at this site. We furthermore generated an S2’ site mutant, SARS2-S2’-AA, by exchanging K814 and R815 to alanine. The S2’ site was shown to be important for proteolytic priming in SARS1-S and is highly conserved among coronavirus spikes (27). We therefore suspected that this site is also important for proteolytic processing of SARS2-S. Clearly detectable bands of lower molecular weight, indicative of proteolytic processing, were only observed with wt SARS2-S (Fig. 1 B). As expected, the S1/S2-mutant exhibited no processing at the S1/S2 site indicated by a missing S2 fragment in Western Blot from transfected 293T lysate (Fig. 1 B). This is similar to SARS1-S which has no furin cleavage site at this position. As not all mutants might be efficiently expressed at the cell surface, we performed cell surface staining with a COVID-19 convalescent serum followed by flow cytometry (Fig. 1 C, middle column group), which revealed detectable but strongly reduced cell surface expression of the SARS2-S2’-AA mutant, as well as reduced ACE2 binding when the same assay was performed with an ACE2-Fc fusion protein (Fig. 1 C, right column group). SARS1-S was only weakly recognized by the COVID-19 convalescent serum. RRV gHΔ21-27-Fc, an Fc fusion protein of RRV gH that lacks any detectable receptor interactions (28), served as control.

**Figure 1:**
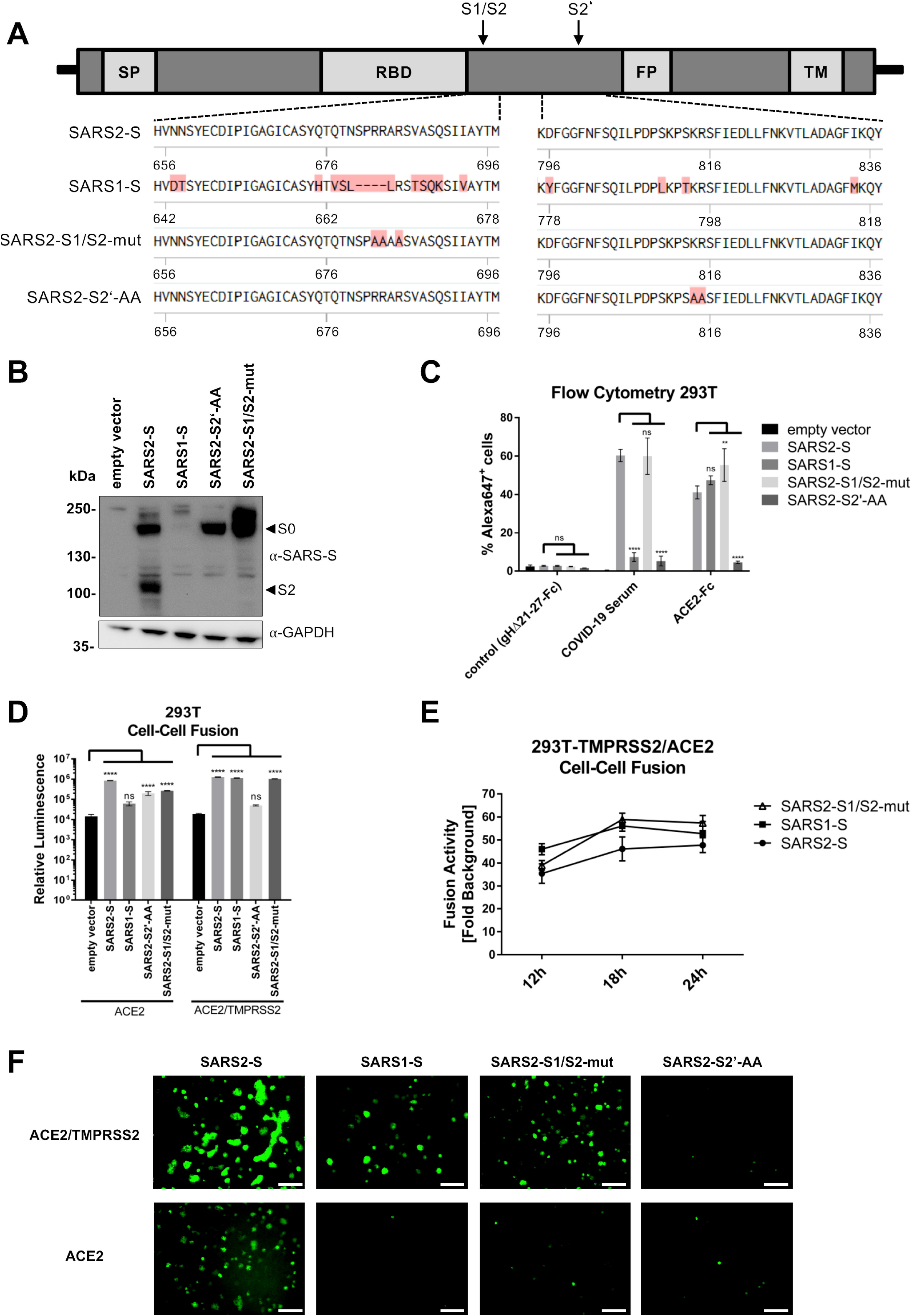
SARS2-S mediates robust fusion activity in the presence of ACE2 or ACE2 and TMPRSS2 on target cells, and ablation of the S1/S2 or S2’ proteolytic cleavage site affects fusion activity differently. **A** Schematic illustration of the coronavirus spike protein showing the signal peptide (SP), the receptor binding domain (RBD), the fusion peptide (FP), the transmembrane domain (TM), the S1/S2 cleavage site (S1/S2) and the S2 cleavage site (S2’), together with amino acid sequence alignments of the spike proteins of SARS-CoV-2, SARS-CoV, and the SARS2-S cleavage site mutants analyzed in this study (not exactly drawn to scale). **B** Expression of spike variants in 293T cells. The unprocessed spike (S0) and the S1/S2-site processed (S2) spike are indicated by arrows. The expression of GAPDH served as loading control. **C** Cell surface expression and ACE2 binding. Cell surface expression as measured by antibody binding from a COVID-19 convalescent serum and binding of soluble ACE2-Fc to 293T cells expressing the indicated spike proteins was determined via flow cytometry analysis and detection with an Alexa647-coupled secondary antibody to human IgG. The percentages of Alexa647-positive cells are shown. Error bars represent the standard deviation of three independent experiments. **D** Cell-cell fusion assay. Effector cells (293T transfected with either empty vector or expression plasmids for the indicated spike variants and Vp16-Gal4 transactivator) were co-cultured with target cells (293T transfected with empty vector or ACE2/TMPRSS2 expression plasmids and Gal4-TurboGFP-Luc reporter plasmid). After 24h luciferase activity was measured. The data shows averaged relative luminescence units, error bars represent the standard deviations of one representative experiment performed in triplicates. **E** Experiment as shown in **D**, except that only ACE2/TMPRSS2 target cells were analyzed. After 12h, 18h and 24h luciferase activity was measured. The data shows averaged fusion activity normalized to empty vector transfected effector cells, error bars represent the standard deviations of one representative experiment performed in triplicates. **F** Representative GFP fluorescence microscopy images of a cell-cell fusion assay with ACE2 and ACE2/TMPRSS2 expressing target cells and effector cells expressing the indicated spike variants (200 μm scale bar). Statistical significance in **C** and **D** was determined by Two-Way ANOVA, p-values were corrected for multiple comparisons by Sidak’s method (p>0.05, ns; p≤0.05, *; p≤0.01, **; p≤0.001, ***; p≤0.0001, ****).

In order to study spike-mediated cell-cell fusion, we established a quantitative reporter gene assay. We chose 293T cells as effector cells, i.e. the cell expressing the viral glycoproteins, because i) 293T exhibit high transfection efficiency and protein expression and ii) 293T can be lifted without trypsinization. We resorted to a system that is also used for two-hybrid screenings, using a VP16-Gal4 transcription factor in one cell and a Gal4 response element-driven reporter construct in the other cell, which results in strong transactivation and reporter gene expression after cell-cell fusion. We transfected 293T target cells with ACE2 and TMPRSS2 expression plasmids and a Gal4 response element driven TurboGFP-luciferase reporter plasmid (Gal4-TurboGFP-Luc) and effector cells with spike expression constructs, as well as with a plasmid encoding the Gal4 DNA binding domain fused to the VP16 transactivator. Apparent expression levels of SARS1-S as assayed by Western blot were lower than those of SARS2-S (Fig. 1 B), but this may be owed to different glycosylation, proteolytic cleavage and transfer or detection and was not reflected in its surface expression as measured by ACE2-binding (Fig. 1 C) and its fusion activity (Fig. 1 D). We found that when only ACE2 was overexpressed (Fig. 1 D, left), all SARS2-S constructs exhibited fusion activity that was statistically different from background. SARS1-S had visible activity but that did not remain significant after correction for multiple comparisons. On 293T cells that were co-transfected with ACE2/TMPRSS2 expression constructs, all spike variants exhibited fusion activity significantly over background (Fig. 1 D, right), except for the SARS2-S2’-AA mutant, which exhibited visible but not statistically significant activity. We chose a logarithmic scale in Fig. 1 D for an initial overview of the considerably different fusion activities and how they relate to background activity. Testing activity in a time-lapse experiment, we observed that luciferase activity was increasing up to 18h for SARS1-S and SARS2-S1/S2-mut and possibly 24h for SARS2-S (Fig. 1 E). Also, activity between SARS1-S, SARS2-S, and SARS2-S1/S2-mut was not meaningfully different at any timepoint. Activity is shown on a linear scale here, which allows for discrimination of smaller differences and which we used from here on.

### The S1/S2 site is critical for syncytium size

Our results demonstrated mostly normal fusion activity of the S1/S2 mutant in our system when TMPRSS2 was present. Therefore, we wanted to address how mutation of the S1/S2 site translates into syncytium formation in our system, as several reports clearly demonstrated that the S1/S2 site is important for this process (24, 26). It should be noted that initial cell-cell fusion and syncytium formation may not necessarily be the exact same thing. After the initial fusion event, all factors that were originally present in separate cells, i.e. viral glycoprotein, receptor, and activating proteases are then together in a single syncytial cell and can interact directly upon co-expression. As our reporter also encodes a TurboGFP that is fused to firefly luciferase, syncytium formation can be conveniently visualized. Under the microscope, we indeed observed that in the presence of ACE2 and TMPRSS2 the S1/S2 mutant formed small but numerous syncytia, while wt SARS2-S formed larger syncytia (Fig. 1 F). Luciferase reporter activity was comparable. Formation of extended syncytia is obviously a quality that our luciferase reporter does not capture, and interestingly, this is not a matter of the timing of the measurement (Fig. 1 E), as even at earlier time points, the luciferase activity between SARS2-S wt and the S1/S2 mutant as well as SARS1-S were similar. We conclude that our luciferase assay measures primarily the initial fusion between effector and target cell and not the formation of extended syncytia.

### SARS2-S-mediated cell-cell fusion is dependent on ACE2 receptor expression and is less restricted by TMPRSS2-mediated activation *in trans* than SARS1-S-mediated fusion

As we found SARS2-S capable of fusing 293T cells efficiently when ACE2 was expressed without TMPRSS2, while SARS1-S was only fully fusogenic in the presence of TMPRSS2, we decided to analyze SARS2-S, SARS1-S and SARS2-S1/S2-mut as well as the SARS2-S2’-AA mutant in the context of different ACE2 and TMPRSS2 expression levels (Fig. 2). In this setting, we again observed robust fusion activity of SARS2-S that was essentially unaltered by different levels of TMPRSS2 but required the presence of ACE2 (Fig. 2 A). SARS1-S on the other hand exhibited high activity under all conditions with TMPRSS2 present, whether ACE2 was recombinantly expressed or not. Activity of the SARS2-S2’-AA mutant was low under all conditions but was highest under the condition with maximal ACE2 expression and not responsive to changes in TMPRSS2 levels. SARS2-S1/S2-mut exhibited an interesting behavior in that it exhibited reduced fusion activity when either ACE2 or TMPRSS2 were absent but was fully fusion competent in all conditions in between, with probably a slight trend towards highest activity with comparatively low TMPRSS2 levels, similar in that respect to SARS1-S. The respective protein levels as present at the end of the co-culture are shown in Fig. 2 B. We labelled the fully processed S2 fragment with an asterisk, as the exact nature of this fragment can’t be deduced with full confidence from its apparent molecular size, even if it could be the so-called S2’ fragment after cleavage at this site. Interestingly, the S0 and S2 fragments of SARS2-S are visibly processed to a large degree into smaller fragments under conditions that allow for high fusion activity. We decided to continue with transfecting equal amounts of ACE2 and TMPRSS2 expression plasmids.

**Figure 2:**
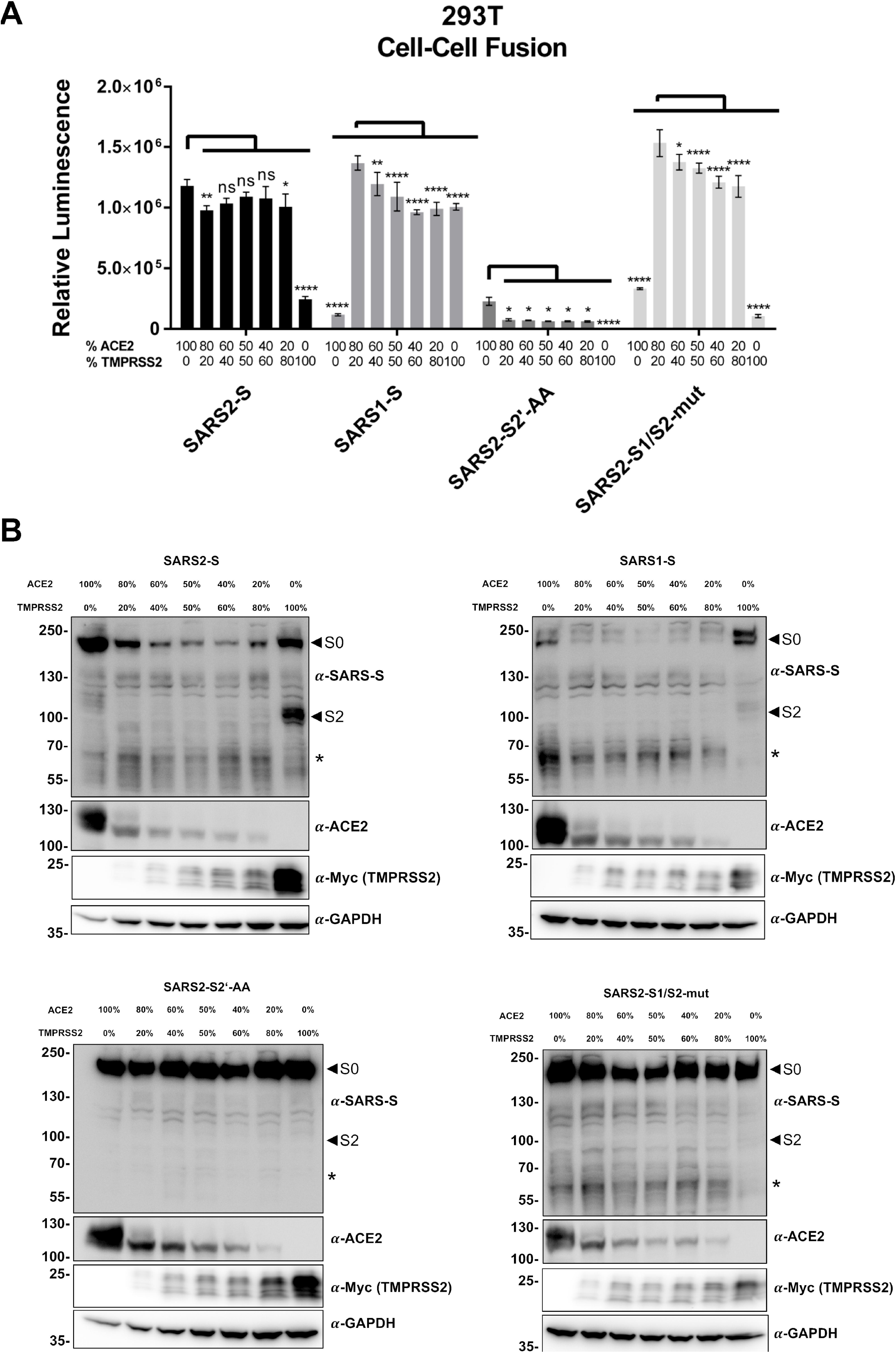
SARS2-S-mediated cell-cell fusion depends on ACE2 receptor expression whereas SARS1-S-mediated fusion depends on TMPRSS2 activity in 293T cells. **A** Cell-cell fusion assay. Effector cells (293T transfected with either empty vector or expression plasmids for the indicated spike variants and Vp16-Gal4 transactivator) were co-cultured together with target cells (293T transfected with ACE2 or TMPRSS2 expression plasmids at the indicated ratios and Gal4-TurboGFP-Luc reporter plasmid). After 24h luciferase activity was measured. The data shows averaged relative luminescence units, error bars represent the standard deviations of one representative experiment performed in triplicates. Comparisons were made against the condition with maximum activation using Two-Way ANOVA, p-values were corrected for multiple comparisons by Sidak’s method (p>0.05, ns; p≤0.05, *; p≤0.01, **; p≤0.001, ***; p≤0.0001, ****). **B** The expression of proteins in target cells and effector cells after co-cultivation was analyzed by Western blot from lysates harvested for determination of luciferase activity in **A**. The unprocessed spike (S0) and the S1/S2-site processed spike (S2) are indicated by arrows. An additional cleavage product marked with an asterisk was observed. The predominant, processed low molecular weight TMPRSS2 fragment is shown. The expression of GAPDH served as loading control. One representative Western blot is shown.

### Differential effect of the TMPRSS2 inhibitors Camostat, Bromhexine, and the Bromhexine metabolite Ambroxol on SARS1-S- and SARS2-S-mediated fusion

For a comprehensive analysis, we measured fusion with target cells that were co-transfected with ACE2 and TMPRSS2 expression plasmids, in addition to cells transfected with either ACE2 or TMPRSS2 expression plasmid alone. As fusion effectors, SARS1-S, SARS2-S as well as SARS2-S1/S2-mut and SARS2-S2’-AA were included. To test the effects of TMPRSS2 inhibition by small molecules on the activation of wt SARS2-S and the two mutants as well as SARS1-S, we incubated the different target cells with Bromhexine, reportedly a specific inhibitor of TMPRSS2 (15), the chemically related compound Ambroxol, or Camostat, an irreversible inhibitor of TMPRSS2 and many serine proteases in general (29, 30), at 50 μM (Fig. 3 A). We chose this high concentration, which is most likely outside of any therapeutic range except for Ambroxol, as overexpression of TMPRSS2 may shift the EC50 considerably upwards.

**Figure 3:**
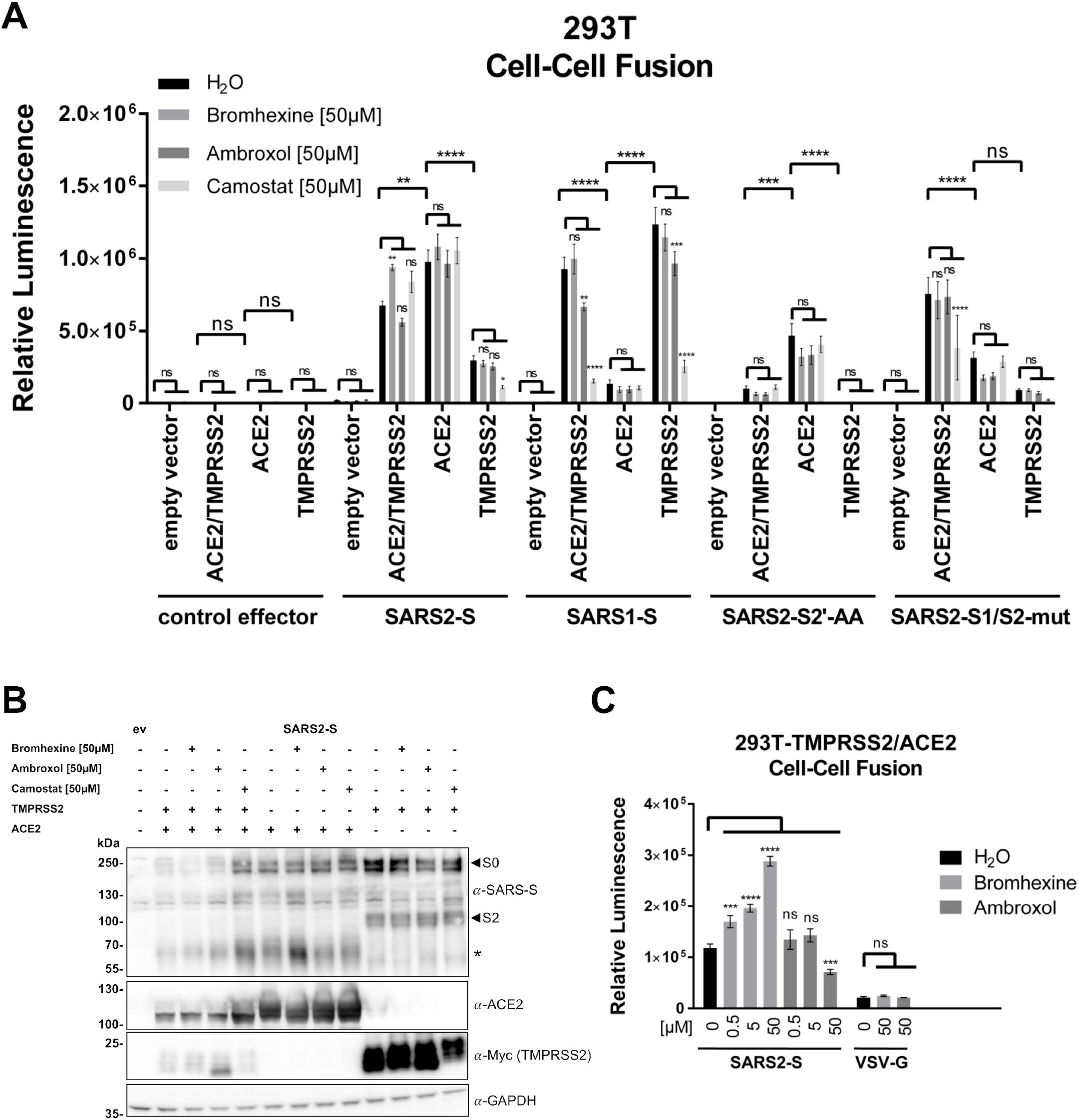
SARS2-S mediated cell-cell fusion of 293T cells is enhanced by Bromhexine in the presence of TMPRSS2. **A** Cell-cell fusion assay. Effector cells (293T transfected with either empty vector or expression plasmids for the indicated spike variants together with Vp16-Gal4 expression plasmid) were added to target cells (293T transfected with empty vector, expression plasmids for ACE2, TMPRSS2 alone or in combination and Gal4-TurboGFP-Luc reporter plasmid), which had been pre-incubated for 30 min with Bromhexine, Ambroxol or Camostat. After addition of effector cells, effector und target cells were co-cultured in the presence of the respective inhibitors at 50μM. After 24h luciferase activity was measured. The data shows averaged relative luminescence units and the error bars represent the standard error of the mean of four independent experiments, each performed in triplicates. Statistical significance was determined by Two-Way ANOVA, p-values were corrected for multiple comparisons by Sidak’s method (p>0.05, ns; p≤0.05, *; p≤0.01, **; p≤0.001, ***; p≤0.0001, ****). For the comparison between inhibitor treatments, the three comparisons within each family were corrected for. The p-values for comparisons between different H_2_O (control) treated target cell populations were corrected for multiple comparison of each target cell and effector cell combination in the inhibitor group (in total 190 possible comparisons). **B** The expression of proteins in treated target cells and effector cells after co-cultivation was analyzed by Western blot from lysates harvested for determination of luciferase activity shown in **A**. The unprocessed spike (S0) and the S1/S2-site processed (S2) spike are indicated by arrows. An additional cleavage product marked with an asterisk was observed. The predominant, processed low molecular weight TMPRSS2 fragment is shown. The expression of GAPDH served as loading control. One representative Western blot is shown. ev = empty vector. **C** Cell-cell fusion assay. Effector cells (293T cells transfected with the indicated glycoprotein expression plasmids and Gal4-Luc reporter plasmid) were co-cultured with target cells (293T transfected with ACE2 and TMPRSS2 expression plasmids and Vp16-Gal4 expression plasmid) that were pre-incubated for 30 min with Bromhexine or Ambroxol. After addition of effector cells, effector und target cells were co-cultured with inhibitors at indicated concentrations, and luciferase activity of cell lysates was measured after 48h. Data shows averaged relative luminescence units of one experiment performed in triplicates, error bars represent the standard deviation.

As observed before (Fig. 1 D), in the presence of ACE2 and TMPRSS2, both SARS1-S and SARS2-S exhibited strong fusion activity, as did SARS2-S1/S2-mut. SARS2-S2’-AA on the other hand was strongly impaired under these conditions.

ACE2 expression alone was sufficient for induction of high fusion activity of SARS2-S but induced only moderate activity of SARS1-S. Levels of ACE2 expression were higher in single-transfected cells (Fig. 3 B). This observation is compatible with data from the literature stating that ACE2 is cleaved by TMPRSS2 (10), which conceivably reduces detection by Western blot, in addition to potential competition effects between expression plasmids.

Nevertheless, SARS2-S-driven fusion was clearly not limited by TMPRSS2 expression, and reached highest activity when only ACE2 was expressed. The S1/S2 cleavage site mutant of SARS2-S on the other hand exhibited reduced activation in the presence of ACE2 without additional TMPRSS2 activity, whereas the SARS-S2’-AA mutant exhibited again low but detectable fusion activity when ACE2 was overexpressed. Overexpression of TMPRSS2 did not increase fusion activity of SARS-S2’-AA. Conversely, SARS1-S-driven fusion was clearly more enhanced by overexpression of TMPRSS2 than by overexpression of ACE2, reaching high activity under conditions where only TMPRSS2 was recombinantly expressed, and was only weakly activated by ACE2 expression in the absence of recombinant TMPRSS2 expression (Fig. 2 A, 3 A). We observed that cell-cell fusion by SARS1-S and SARS2-S was not inhibited by Bromhexine, and only SARS1-S activity was slightly inhibited by Ambroxol in the presence of TMPRSS2.

Surprisingly, we observed an induction of SARS2-S fusion activity in the presence of Bromhexine, significantly so when ACE2 and TMPRSS2 were co-expressed. Camostat did not reduce SARS2-S-mediated fusion in this setting unless TMPRSS2 was overexpressed without ACE2. However, both SARS2-S1/S2-mut and even more pronouncedly SARS1-S exhibited a significantly reduced fusion activity in the presence of Camostat. The strong induction of SARS1-S-mediated fusion by TMPRSS2 was clearly reversed by Camostat but not by Bromhexine. Notably, Camostat did not exert any inhibitory effect on the remaining fusion activity of the SARS2-S2’-AA mutant, nor did TMPRSS2 expression induce activity of this mutant, compatible with the S2’ site being the primary target of TMPRSS2-mediated activation *in trans*.

The results were also mirrored by Western blot analysis (Fig. 3 B) of SARS2-S under the same conditions, if generation of the fully processed S2 fragment is analyzed, which we labelled with an asterisk. Generation of this fragment was clearly visible under all conditions that allowed for high fusion activity, e.g. when ACE2 was present, less so with TMPRSS2 alone. Interestingly, addition of Camostat increased the detectable amount of ACE2, probably explaining the slight trend towards higher activity in its presence. Further, Ambroxol reproducibly induced the generation of an atypical TMPRSS2 autoproteolytic fragment, which may hint at some sort of modulating activity of Ambroxol towards TMPRSS2 (Fig. 3 B, fourth lane).

Taken together, we observed robust SARS2-S-mediated cell-cell fusion with ACE2 overexpressing cells that was not dependent on exogenous TMPRSS2 expression and that was not inhibited by Bromhexine. Instead, fusion was enhanced by Bromhexine. Cell-cell fusion mediated by SARS2-S was clearly not at all or to a much lesser degree restricted by serine protease activity on target cells than fusion by SARS1-S. Interestingly, Ambroxol exhibited some activity against TMPRSS2-mediated activation of SARS1-S.

### Bromhexine enhances SARS2-S-mediated fusion in the presence of TMPRSS2

To further explore the paradoxical effect of the putative TMPRSS2 inhibitor Bromhexine on fusion activity, we performed fusion reactions in the presence of Bromhexine and Ambroxol at different concentrations (Fig. 3 C). In order to eliminate potential systematic errors, we deviated from our previous protocol, co-cultured for 48h instead of 24h, and co-transfected the reporter plasmid into the effector instead of the target cells, this time using a different luciferase reporter without TurboGFP. We again did not observe inhibition by Bromhexine, but a dose-dependent enhancement. Ambroxol treatment on the other hand did not lead to a similar enhancement, but to a slight decrease in activity at 50μM. As a control fusion protein that works with practically any cell type, we included VSV-G. While VSV-G is physiologically pH-activated for full fusion activity (31), it reportedly exhibits considerable activity without pH priming (32, 33). VSV-G-mediated fusion activity was not increased by Bromhexine.

### SARS2-S-mediated cell-cell fusion is sensitive to inhibition of matrix metalloproteases

The robust cell-cell-fusion that we observed with SARS2-S in the absence of TMPRSS2 activity should most likely be triggered by proteolytic processing, if the mechanism is analogous to what was observed for SARS-CoV (8, 34). Therefore, we tested the effects of different protease inhibitors on SARS2-S-mediated fusion of ACE2 expressing 293T cells without exogenous TMPRSS2 activity. As we wanted to exclude the possibility that pre-activation on the producer cells could play a role, we tested the inhibitors both in the co-culture (Fig. 4 A), and with pre-incubation of both effector and target cells (Fig. 4 B). Values were normalized to the respective solvent control for better comparison. We observed some inhibitory effect on SARS2-S and SARS1-S fusion activity by the broadband serine protease inhibitor AEBSF, and by a protease inhibitor cocktail whose main ingredients are the serine protease inhibitors AEBSF and Aprotinin, and the cysteine protease inhibitors E64 and Leupeptin. The S1/S2 cleavage site mutant was not sensitive to this inhibitor cocktail, suggestive of action at this site in the SARS-CoV-2 wildtype spike. These effects were more pronounced and significant with pre-incubation of the effector cells (Fig. 4B), in particular for the SARS2-S2’-AA mutant. Interestingly, the inhibitor cocktail almost completely abrogated the remaining fusion activity of SARS1-S. The furin inhibitor CMK did not significantly inhibit any of the spikes except for the S2’ mutant (Fig. 4 A, B). This was somewhat surprising for us, but it may reflect the fact that proteases other than furin can cleave at the S1/S2 site (35), which may in turn partially obviate furin cleavage in our system. We also tested EDTA/EGTA, Bromhexine, Ambroxol, and Camostat, which as expected had no effect in this TMPRSS2-free system. EDTA/EGTA had a mild impact on SARS1-S fusion activity with and without pre-incubation (Figs. 4 A, B). Bromhexine and Ambroxol exhibited an interesting behavior in this assay: We observed inhibitory activity of Bromhexine and Ambroxol towards SARS1-S and the SARS2-S1/S2-mut and SARS2-S2’-AA mutants in this TMPRSS2-free cell system, suggesting that these substances somehow interact with the spike proteins or ACE2. Luciferase activity of control cells, which were transfected with both the Gal4-reponse element-driven reporter and the Gal4 transactivator constructs, was only mildly affected by AEBSF, the inhibitor cocktail, EDTA/EGTA and Bromhexine, not by the other substances (Fig. 4 C). In particular, any reductions observed with Ambroxol can’t be explained by non-specific effects on the luciferase reporter system and most likely represent real inhibitory activity against SARS1-S-mediated fusion activity and fusion mediated by the two SARS2-S cleavage site mutants.

**Figure 4:**
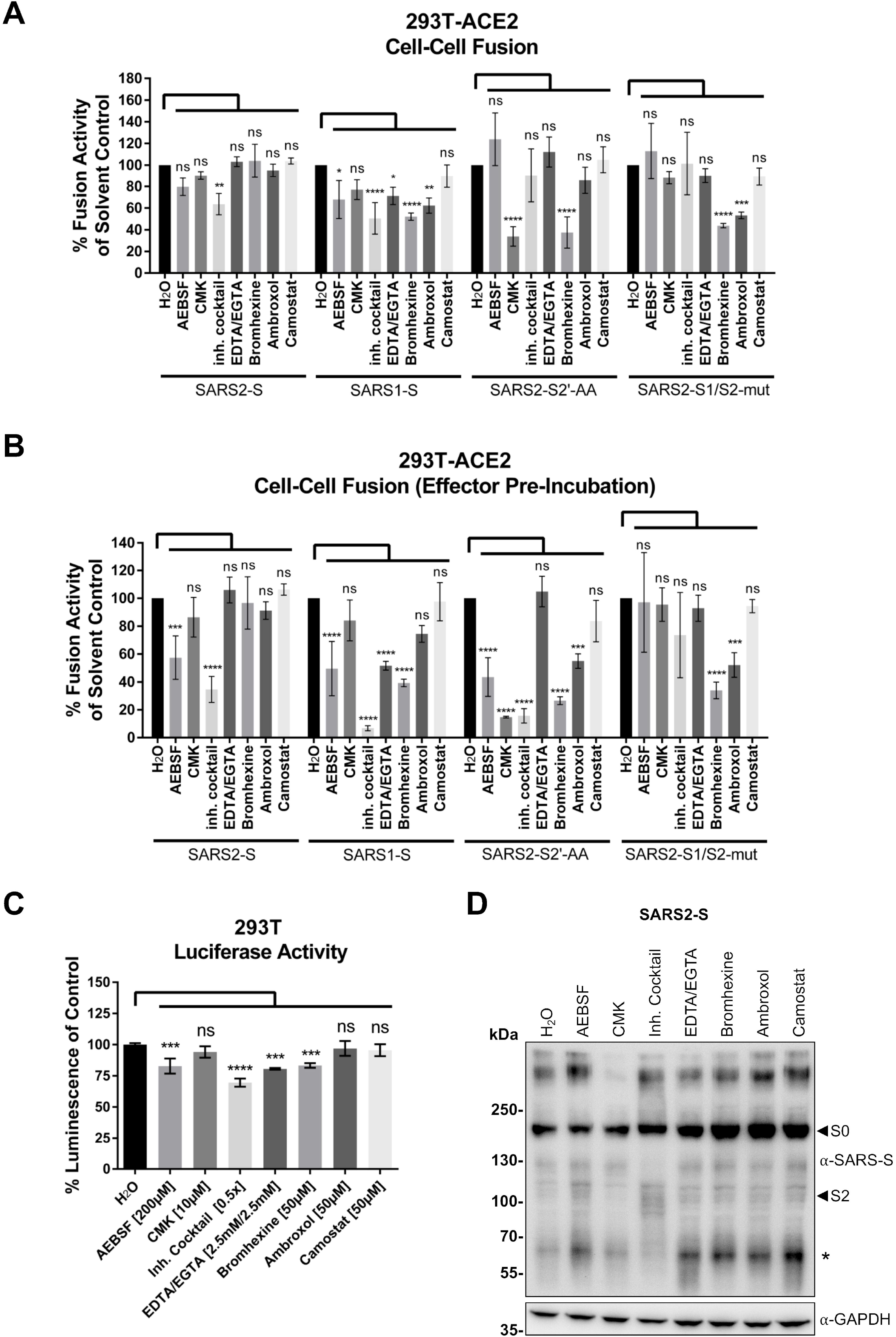
Sensitivity of SARS2-S-mediated 293T cell-cell fusion to different inhibitors. **A** Cell-cell fusion assay. Effector cells (293T transfected with expression plasmids for the indicated spike variants together with Vp16-Gal4 expression plasmid) were added to target cells (293T transfected with expression plasmids for ACE2 and Gal4-TurboGFP-Luc reporter plasmid), which had been pre-incubated for 30 min with twice the final concentration of AEBSF (200μM), furin inhibitor CMK (10μM), proteinase inhibitor cocktail, EDTA/EGTA (2.5mM each), Bromhexine (50μM), Ambroxol (50μM) and Camostat (50μM). After addition of effector cells, effector und target cells were co-cultured in the presence of the respective inhibitors. After 24h luciferase activity was measured. The data shows values normalized to solvent treatment which was set to 100% and the error bars represent the standard deviation of three independent experiments, each performed in triplicates. **B** Cell-cell-fusion assay as shown in **A**, except that effector cells were pre-incubated with indicated inhibitors for 18h before co-culture with target cells. The target cells were pre-incubated with indicated inhibitors for 30min before addition of effector cells. After 24h luciferase activity was measured. The data shows values normalized to solvent treatment which was set to 100% and the error bars represent the standard deviation of three independent experiments, each performed in triplicates. **C** 293T cells transfected with Vp16-Gal4 and Gal4-TurboGFP-Luc reporter expression plasmids and were incubated with inhibitors as described in **A**. After 24h luciferase activity was measured. The data shows values normalized to solvent treatment which was set to 100%, error bars represent the standard deviations of one representative experiment performed in triplicates. **D** The expression of proteins in treated target cells and effector cells after co-cultivation was analyzed by Western blot from lysates harvested for determination of luciferase activity shown in **A**. The unprocessed spike (S0) and the S1/S2-site processed (S2) spike are indicated by arrows. An additional cleavage product marked with an asterisk was observed. The predominant, processed low molecular weight TMPRSS2 fragment is shown. The expression of GAPDH served as loading control. One representative Western blot is shown.

Western blot analysis suggested that the protease inhibitor cocktail may have had a somewhat stabilizing effect on the S2 intermediate form of SARS2-S (Fig. 4 D), which resulted in less processing into the putative S2’ form (marked by an asterisk). CMK both reduced “smear” at higher molecular weight, which likely represents glycosylation variants, and reduced abundance of the S2 proteolytic product, that should be generated through cleavage at the polybasic cleavage site, compatible with furin inhibition. As none of the tested inhibitors resulted in meaningful reduction of fusion activity of wt SARS2-S that could not also be explained by toxicity, we decided to test a more potent inhibitor of metalloproteases than EDTA/EGTA, whose maximum concentration is limited by its effects on cell adhesion and viability. The EDTA/EGTA concentration that was used by us was most likely too low to meaningfully impact protease activity, in particular as the cell culture medium contains calcium and magnesium. We therefore tested Batimastat, which inhibits matrix metalloproteases (36, 37).

Batimastat indeed inhibited SARS2-S-dependent fusion in the absence of TMPRSS2 in a dose dependent manner (Fig. 5 A). Interestingly, no inhibition was observed in the presence of both ACE2 and TMPRSS2, and in the presence of TMPRSS2 alone unless TMPRSS2 was inhibited by Camostat (Fig. 5 A). Therefore, Batimastat-sensitive metalloproteases cleave SARS2-S to activate cell-cell fusion. This notion is supported by the finding that TMPRSS2 expression can overcome the Batimastat-induced block. Western blot analysis of the fusion reactions indicated that Batimastat probably induced a subtle change in the migration pattern of the SARS2-S S2 fragment in the presence of ACE2 but without TMPRSS2 (Fig. 5 B). We next decided to test the effect of Batimastat on the fusion activity of the S1/S2 mutant and the S2’-AA mutant under conditions of ACE2 overexpression without TMPRSS2 (Fig. 5 C, left). Both mutants were inhibited by Batimastat, indicating that matrix metalloproteases can cleave irrespective of an intact S1/S2 or S2’ cleavage site, although this does not necessarily rule out an modulating effect in particular by S1/S2 cleavage, as mutation of S1/S2 leads to impaired activity without TMPRSS2. SARS1-S was also slightly affected by Batimastat under these conditions, but at an overall very low activity level (compare Fig. 5 A). Under conditions of ACE2 and TMPRSS2 co-expression, which leads to lower ACE2 levels (compare Figs. 2 B, 3 B, 5 B), SARS2-S1/S2-mut was not impacted by Batimastat unless TMPRSS2 was again inhibited by addition of Camostat (Fig. 5 C, middle), whereas activity of the S2’ mutant was inhibited in the presence of Batimastat alone, strongly suggesting that TMPRSS2 activates via the S2’ site. Under conditions of TMPRSS2 overexpression without ACE2 overexpression (Fig. 5 C, right) Batimastat was again without effect. Results with the SARS2-S2’-AA mutant come with the caveat that this mutant was barely active at all under these conditions (Figs. 2 A, 3 A). In summary, these experiments demonstrate that in the presence of the ACE2 receptor, matrix metalloproteases can efficiently activate SARS2-S for cell-cell fusion.

**Figure 5:**
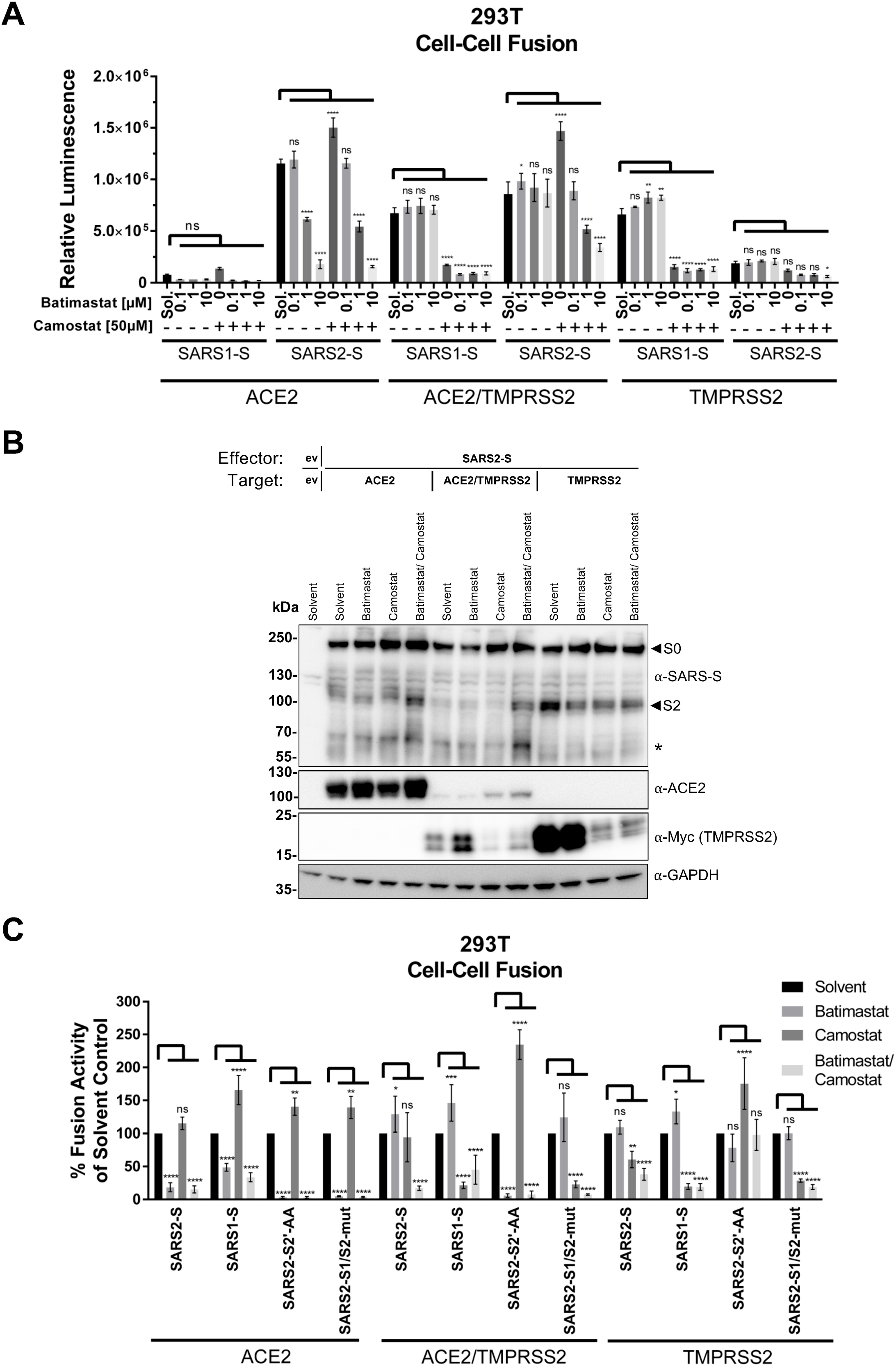
The matrix metalloproteinase inhibitor Batimastat inhibits SARS2-S-mediated cell-cell fusion. **A** Cell-cell fusion assay: Effector cells (293T transfected with expression plasmids for the indicated spike variants together with Vp16-Gal4 expression plasmid) were added to target cells (293T transfected with expression plasmids for ACE2, ACE2/TMPRSS2, TMPRSS2 and Gal4-TurboGFP-Luc reporter plasmid), which had been pre-incubated with Batimastat or Camostat for 30 min at twice the indicated final concentration. After 24h luciferase activity was measured. The data shows averaged relative luminescence units, error bars represent the standard deviations of one representative experiment performed in triplicates. **B** The expression of proteins in treated target cells and effector cells after co-cultivation was analyzed by Western blot from lysates harvested for determination of luciferase activity shown in **A**. The unprocessed Spike (S0) and the S1/S2-site processed (S2) spike are indicated by arrows. An additional cleavage product marked with an asterisk was observed. The predominant, processed low molecular weight TMPRSS2 fragment is shown. The expression of GAPDH served as loading control. One representative Western blot is shown. ev = empty vector. **C** Cell-cell fusion assay. Effector cells (293T transfected with expression plasmids for the indicated spike variants together with Vp16-Gal4 expression plasmid) were added to target cells (293T transfected with expression plasmids for ACE2, ACE2/TMPRSS2, TMPRSS2 and Gal4-TurboGFP-Luc reporter plasmid), which had been pre-incubated for 30 min with Batimastat (10μM) and/or Camostat (50μM) at twice the final concentration. After 24h luciferase activity was measured. The data shows values normalized to solvent treatment which was set to 100% and the error bars represent the standard deviation of three independent experiments, each performed in triplicates. Statistical significance in **A** and **C** was determined by Two-Way ANOVA, p-values were corrected for multiple comparisons by Sidak’s method (p>0.05, ns; p≤0.05, *; p≤0.01, **; p≤0.001, ***; p≤0.0001, ****).

### The SARS2-S S2’ site is the target site for TMPRSS2-mediated proteolytic activation

While our results with the SARS2-S2’-AA mutant were already strongly suggestive of S2’ being the target site for TMPRSS2, this conclusion remained slightly ambiguous in light of the relatively low surface expression and inefficient proteolytic processing of this mutant (Fig. 1 B and C). We therefore set out to generate an S2’ mutant that is still efficiently processed and expressed at the cell surface. We permutated several amino acids to replace the original “KR” (Fig. 1 A) sequence motif and tested fusion activity in the presence of ACE2, TMPRSS2 and ACE2/TMPRSS2. We found that SARS2-S2’-GH and -HH mutants were active in our fusion assay, whereas EE and ES resulted in abrogation of fusion activity, below the levels achieved with the AA mutant (Fig. 6 A). The GH mutant was also processed (Fig. 6 B), efficiently expressed at the cell surface, and exhibited high ACE2 binding capacity (Fig. 6 C). For further experiments, we continued with the SARS2-S2’-GH mutant. Interestingly, when we tested the furin inhibitor CMK for its effects in absence of TMPRSS2, all spike variants were slightly less active, but only the S2’-GH variant was significantly inhibited, suggesting increased dependence on pre-priming by furin in absence of the S2’ site (Fig. 6 D).

**Figure 6:**
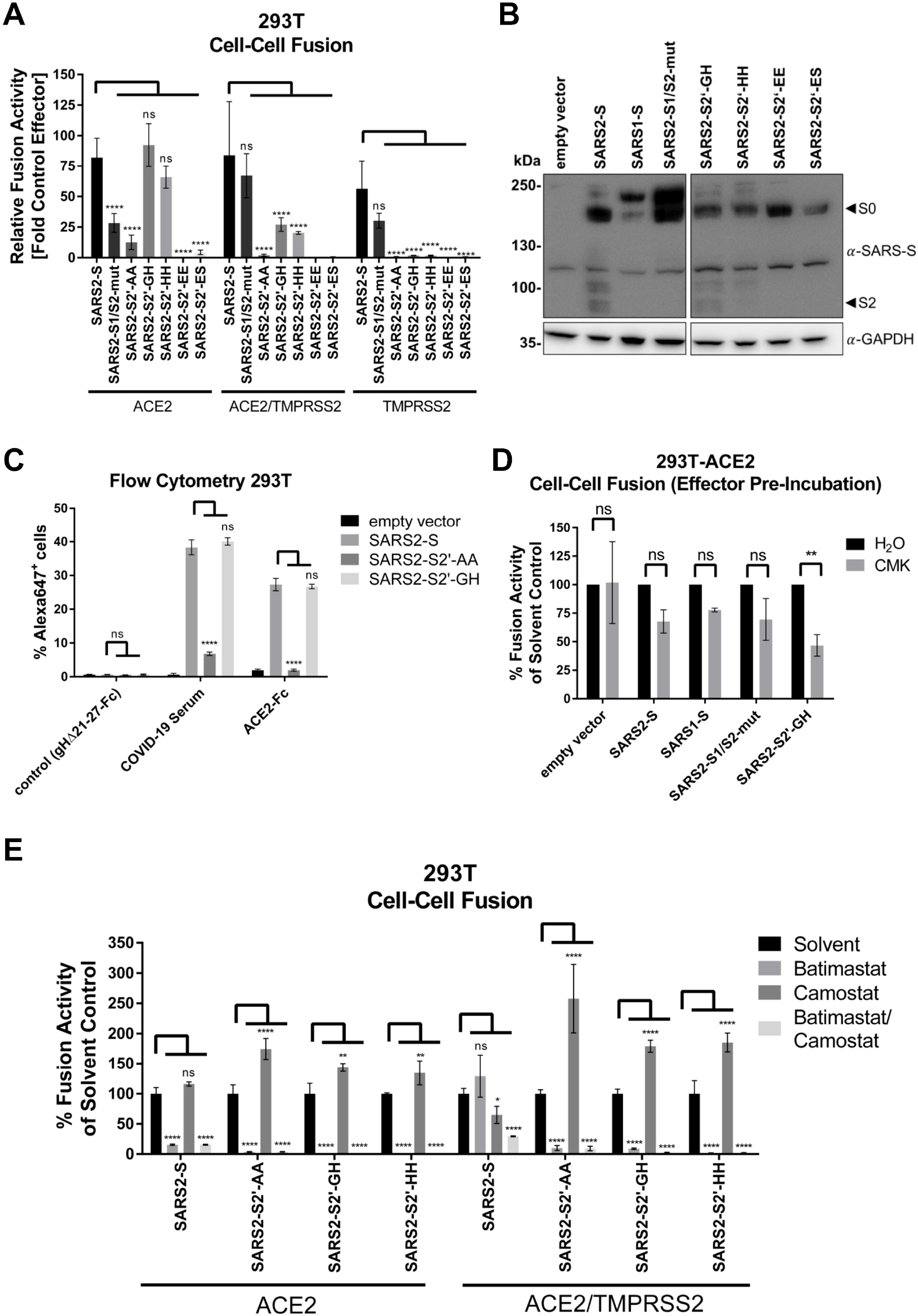
The conserved S2’ site is the site of TMPRSS2-mediated activation of SARS2-S for cell-cell fusion. **A** Cell-cell fusion assay. Effector cells (293T transfected with expression plasmids for the indicated spike variants together with Vp16-Gal4 expression plasmid) were added to target cells (293T transfected with expression plasmids for ACE2, ACE2/TMPRSS2, TMPRSS2 and Gal4-TurboGFP-Luc reporter plasmid). After 24h luciferase activity was measured. The data shows values fold empty vector control and the error bars represent the standard deviation of three independent experiments, each performed in triplicates. **B** Expression of analyzed spike variants in 293T cells. The unprocessed spike (S0) and the S1/S2-site processed spike (S2) are indicated by arrows. The expression of GAPDH served as loading control. **C** Cell surface expression and ACE binding. Cell surface expression and binding of soluble ACE2-Fc by the indicated spike variants was determined by flow cytometry. Analysis was performed as in Fig. 1 C. **D** Cell-cell fusion assay. Effector cells (293T transfected with expression plasmids for the indicated spike variants together with Vp16-Gal4 expression plasmid) were pre-incubated with furin inhibitor CMK (10μM) and after 16h added to target cells (293T transfected with expression plasmids for ACE2 and Gal4-TurboGFP-Luc reporter plasmid), which had been pre-incubated for 30 min with the same inhibitor concentration. After addition of effector cells, effector und target cells were co-cultured in the presence of CMK. After 24h luciferase activity was measured. The data shows values normalized to solvent treatment which was set to 100% and the error bars represent the standard deviation of two independent experiments, each performed in triplicates. **E** Cell-cell fusion assay. Effector cells (293T transfected with expression plasmids for the indicated spike variants together with Vp16-Gal4 expression plasmid) were added to target cells (293T transfected with expression plasmids for ACE2, ACE2/TMPRSS2, TMPRSS2 and Gal4-TurboGFP-Luc reporter plasmid), which had been pre-incubated with Batimastat and/or Camostat for 30 min at twice the final concentration; final concentrations were Batimastat 10μM and/or Camostat 50μM. After 24h luciferase activity was measured. The data shows values normalized to solvent treatment which was set to 100% and the error bars represent the standard deviation of two independent experiments, each performed in triplicates. Statistical significance in **A, C, D** and **E** was determined by Two-Way ANOVA, p-values were corrected for multiple comparisons by Sidak’s method (p>0.05, ns; p≤0.05, *; p≤0.01, **; p≤0.001, ***; p≤0.0001, ****).

Confirming the results of our prior fusion assays with the AA mutant, also the SARS2-S2’-GH and SARS-S2’-HH fusion activity on 293T cells in the presence of only ACE2 was sensitive to Batimastat (Fig. 6 E, left), and on 293T cells expressing ACE2/TMPRSS2 both SARS2-S2’-GH and SARS-S2’-HH were insensitive to Camostat, but again highly sensitive to Batimastat (Fig. 6 E, right). Fusion activity of the S2 mutants was even increased in the presence of Camostat, likely because inhibition of TMPRSS2 increases ACE2 levels, as demonstrated in Fig. 3 B. This unequivocally identifies the S2’ site as the TMPRSS2 target site, and interestingly as the only TMPRSS2 target site, at least for activation of fusion.

### Entry of SARS2-S-pseudotyped lentiviruses is enhanced by TMPRSS2 and is not inhibited by Bromhexine

To compare our findings on cell-cell fusion to spike protein-driven entry, we used lentiviral particles expressing GFP as reporter gene, pseudotyped with SARS2-S. We found that TMPRSS2 expression was clearly required for efficient infection of 293T cells by SARS2-S-pseudotyped particles (Fig. 7 A). ACE2 overexpression alone also enhanced infection, but considerably less efficiently and barely above the detection limit, which may be owed to our lentiviral GFP system. The TMPRSS2-mediated enhancement was reduced by addition of Camostat, but not by addition of Bromhexine or Ambroxol, both of which may even slightly enhance infection in this setting. These observations were corroborated by fluorescence microscopy (Fig. 7 B). As luciferase is more sensitive than GFP as a reporter gene, we switched to luciferase detection (Fig. 7 C). We also included the SARS2-S D614G variant. As previously reported, D614G-driven infection was more efficient (38). It was also strongly enhanced by TMPRSS2, as evidenced by potent Camostat-mediated inhibition. Ambroxol and Bromhexine had no activity in this system, as opposed to Camostat. Batimastat did not alter SARS2-S driven entry. A VSV-G pseudotyped lentivirus was not significantly affected by either substance.

**Figure 7:**
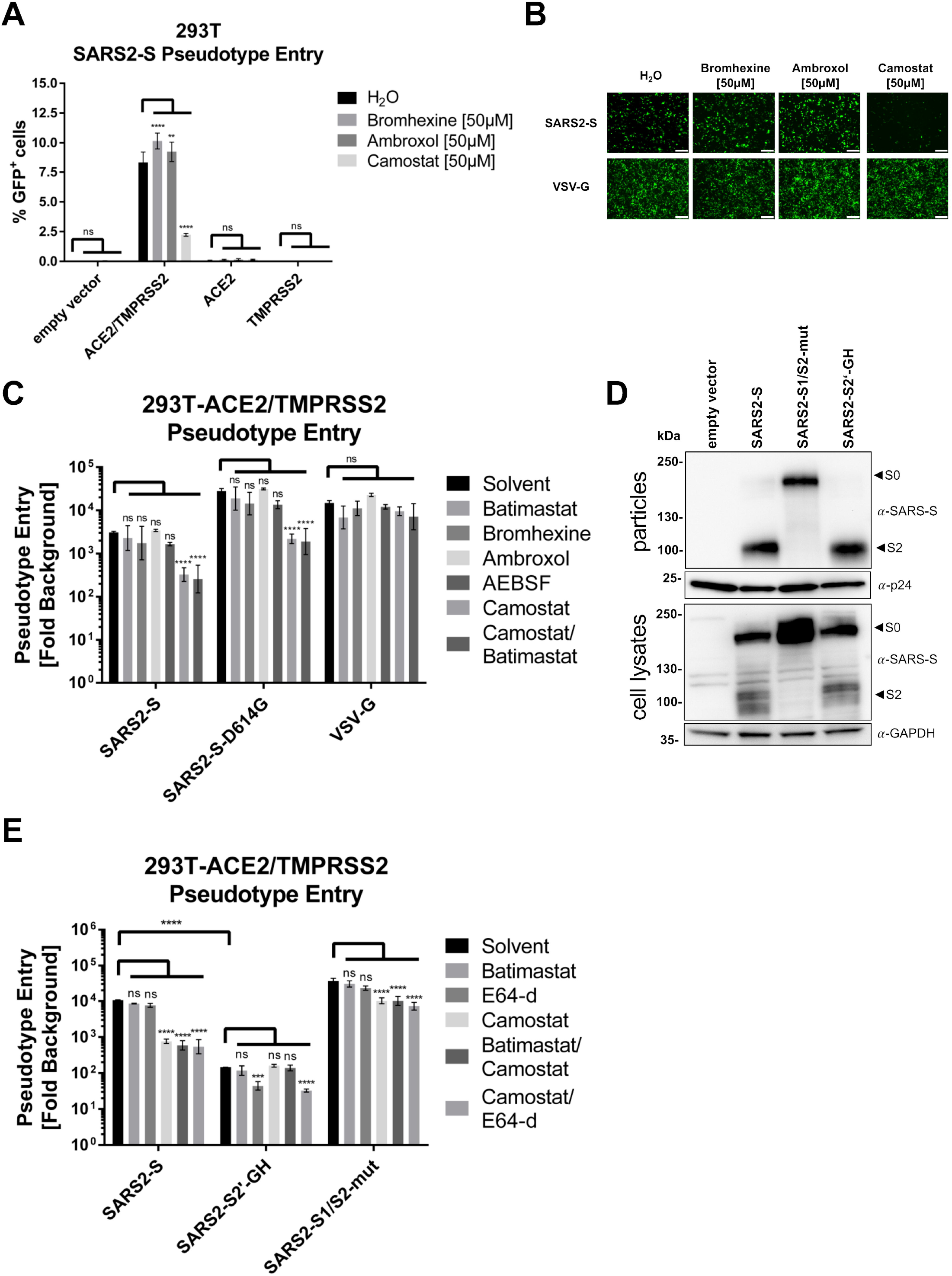
Requirements for the entry of SARS2-S pseudotyped lentiviral particles differ from requirements for SARS2-S-mediated cell-cell fusion. **A** 293T cells transfected with empty vector, ACE2/TMPRSS2, ACE2 or TMPRSS2 were pre-incubated with Bromhexine, Ambroxol or Camostat at the indicated concentration before addition of lentiviral particles pseudotyped with SARS2-S. 48h after transduction the cells were analyzed via flow-cytometry. Data shows averaged percent of GFP-positive cells, error bars represent the standard deviations of one representative experiment performed in triplicates. **B** Micrographs of ACE2 and TMPRSS2 transfected cells that were infected with the respective lentiviral GFP-encoding pseudotype particles. **C** 293T cells transfected with ACE2/TMPRSS2 were pre-incubated with Batimastat (10μM), Bromhexine (50μM), Ambroxol (50μM), AEBSF (200μM), Camostat (50μM) or Batimastat (10μM) in combination with Camostat (50μM) before addition of lentiviral particles pseudotyped with respective glycoprotein. 48h after transduction the cells were lysed and luciferase activity was determined. Data shows fold change over background (bald particles with solvent control), error bars represent the standard deviations of three independent experiments, each performed in triplicates, raw values were log10-transformed before analysis. **D** Western Blot analysis of incorporation of the respective spike variants into lentiviral particles used in **E** and lysate control of transfected 293T cells used for production of lentiviral particles. p24 and GAPDH served as loading control. **E** 293T cells transfected with ACE2/TMPRSS2 were pre-incubated with Batimastat (10μM), E64-d (25μM), Camostat (50μM) or Batimastat (10μM)/ E64-d (10μM) in combination with Camostat (50μM) before addition of lentiviral particles pseudotyped with the respective glycoproteins. 48h after transduction the cells were lysed and luciferase activity was determined. Data shows fold change over background (bald particles with solvent control), error bars represent the standard deviations of two independent experiments, each performed in triplicates, raw values were log10-transformed before analysis. Statistical significance in **A**, **C** and **E** was determined by Two-Way ANOVA, p-values were corrected for multiple comparisons by Sidak’s method method (p>0.05, ns; p≤0.05, *; p≤0.01, **; p≤0.001, ***; p≤0.0001, ****).

### Mutation of the S2’ site uncouples infection from TMPRSS2

Next, we aimed to corroborate our findings regarding the S2’ site as TMPRSS2 target site for cell-cell fusion in pseudoparticle infection. Our SARS2-S2’-GH mutant was efficiently incorporated into lentiviral particles, as was SARS2-S1/S2-mut (Fig. 7 D). Both spike mutants could drive entry into 293T cells expressing ACE2/TMPRSS2, but SARS2-S2’-GH with reduced efficiency and SARS2-S1/S2-mut probably with increased efficiency, although we did not test for the latter (Fig. 7 E). None of the spike variants was inhibited by Batimastat. SARS2-S wt and SARS2-S1/S2-mut were inhibited by Camostat, but not by Batimastat or E64-d, indicating proteolytic activation by TMPRSS2. The S2’ mutant on the other hand was exclusively inhibited by E64-d, indicating that it was refractory to activation by TMPRSS2 and dependent on activation by Cathepsins.

### SARS-CoV-2 is weakly inhibited by Ambroxol on Calu-3 lung cells

As transfected 293T cells express TMPRSS2 at high and possibly variable levels between cells, and allow for at least some entry via endocytosis, weak modulatory effects on ACE2 or TMPRSS2 might be missed in that system. Calu-3 cells express TMPRSS2 to much higher levels than 293T (Fig. 8 A), which are practically negative, but still at endogenous levels. We therefore infected the Calu-3 lung cell line with our lentiviral pseudoparticles (Fig. 8 B). These cells allow for infection by our lentiviral pseudoparticles only at very low levels (not shown). In order to achieve infection at faithfully detectable levels, we used the D614G variant, which exhibited the same sensitivity profile to inhibitors but was about one log more efficient at driving entry (Fig. 7 C). By now, D614G has become the dominant variant globally and is therefore probably also more relevant. As expected, SARS2-S-driven entry was practically abrogated by 50μM Camostat. Bromhexine again had no detectable impact on SARS2-S-driven entry. Ambroxol on the other hand exhibited a weakly inhibitory effect on SARS2-S-driven infection in this system, even if that needs a linear scale for proper visualization. Interestingly, both substances, but Bromhexine more so, affected entry of VSV-G-pseudotyped particles negatively. This is likely owed to the targeting of lysosomal processes by these two substances (39, 40). Finally, we wanted to test whether this small but detectable effect would translate into inhibition of authentic virus. We therefore infected Calu-3 with a clinical isolate of SARS-CoV-2 at low MOI and quantified the viral RNA after 20-24h by RT-qPCR (Fig. 8 C). We chose 5μM and 50μM as concentrations for Ambroxol and Bromhexine, 10μM for Batimastat. For Ambroxol, which is heavily enriched in lung tissue, 50μM might be a clinically attainable concentration. Interestingly, for Ambroxol and Bromhexine, viral RNA copy number trended lower upon treatment, and in a dose-dependent manner as can be observed in the raw Ct values (Fig. 8 C) and after relative quantification (Fig. 8 D). Both, reduction by Ambroxol and by Bromhexine at 50μM was significant, even if inhibition by Ambroxol remained only significant without correction for multiple comparisons, which is appropriate in light of a dose response (ANOVA with post test for linear trend in Ct values at 0μM, 5μM, and 50μM, significant for Ambroxol and Bromhexine). 10μM Batimastat (compared to the DMSO solvent) had no significant effect on SARS-CoV-2 mRNA level, even if DMSO alone had quite some impact compared to water, most likely due to the high concentration needed, which was 1%. In a cell viability assay with Calu-3 using dilutions of commercial over-the-counter cough thinners, neither Bromhexine nor Ambroxol exhibited significant effects up to 10μM (Fig. 8E). We used the cough thinners as an alternative source of Ambroxol and Bromhexine for some control experiments, which were not included in this manuscript, to control for specificity of the observed effects and their independence from the source of the two substances. Bromhexine but not Ambroxol clearly impacted cell viability at 100μM, which is compatible with our observations on 293T (Fig. 4C), although it should be noted that toxicity of Bromhexine may have been overestimated in Fig. 8 E due to non-active ingredients of the cough thinner.

**Figure 8:**
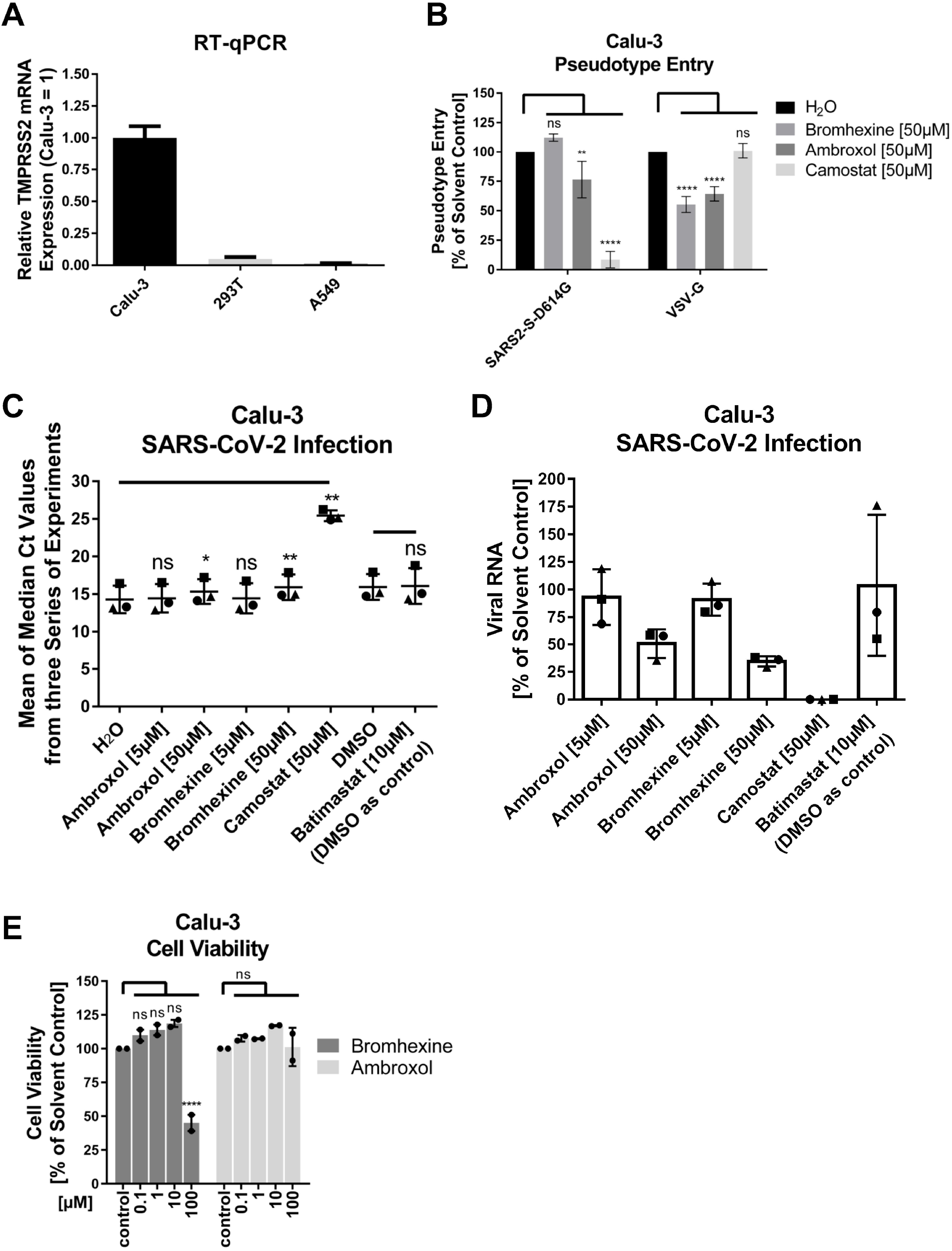
SARS-CoV-2 is weakly inhibited by Bromhexine and Ambroxol on Calu-3 cells. **A** RT-qPCR analysis of TMPRSS2 expression. Fold TMPRSS2 mRNA expression in Calu-3 cells, 293T cells, and A549 cells was measured by RT-qPCR using the ΔΔCt method. The -RT control for the GAPDH mRNA was not negative as expected but the contamination was considered irrelevant as its Ct was more than 19 cycles over the value of the sample, representing a contamination of less than 0.01%. Error bars represent the upper error bound calculated from the sum of the SDs of the ΔCt values for each cell line. **B** Calu-3 cells were infected with lentiviral particles encoding a TurboGFP-luciferase reporter gene pseudotyped with SARS2-S in the presence of 50μM Bromhexine, Ambroxol or Camostat. 48h after transduction the cells were lysed and luciferase activity was determined. The data shows values normalized to solvent treatment, which was set to 100%, and the error bars represent the standard deviation of three independent experiments, each performed in triplicates. **C** Viral RNA load. Calu-3 cells were infected with SARS-CoV-2 in the presence of Bromhexine, Ambroxol, Camostat, or Batimastat at the indicated concentrations. Viral RNA was quantified by RT-qPCR 24h (experiment 1 and 2) and 20h (experiment 3) post infection. The median Ct values of three experiments (each experiment was performed in biological triplicates) are plotted (experiment 1: dots, experiment 2: triangles, experiment 3: squares), and the mean was determined. Significant differences to solvent controls are indicated by asterisks. Significance was determined using repeated measures one way ANOVA and Fisher’s LSD without correcting for multiple comparisons. Differences were also significant using two way ANOVA without correction for multiple comparisons and all available data, but use of the median from each experiment reduced variance. All samples were compared to water except for Batimastat, which was compared to DMSO. **D** Relative viral RNA expression. Using the median Ct values from each experiment series as above and the experimentally determined PCR efficiency, the amount of viral RNA was calculated as percent of solvent control for each inhibitor. **E** Cell viability. Cell viability of Calu-3 was determined after culture in the presence of the indicated compounds in two independent assays, each performed in biological triplicates. Statistical significance was determined using Two-Way ANOVA.

## DISCUSSION

We have established a two-hybrid-based protocol for measuring spike-mediated cell-cell fusion that allows for the quantitation of cell-cell fusion by luciferase activity and visualization of syncytia by GFP fluorescence. Our finding that SARS1-S-mediated and SARS2-S-mediated fusion activity is activated by the ACE2 receptor is in accordance with published data (11), whereas our finding that SARS2-S-mediated cell-cell fusion is relatively more restricted by ACE2 expression and less by proteolytic activation than SARS1-S-mediated fusion is novel. This is because SARS-CoV-2 can efficiently utilize metalloproteases for activation of cell-cell fusion. Further, we have faithfully established the S2’ site of SARS2-S as the target for TMPRSS2-mediated activation through generation of a mutant that is defective for TMPRSS2 activation but otherwise fully functional.

In our system, TMPRSS2 co-expression on ACE2 expressing target cells was not required for SARS2-S-mediated fusion of ACE2 overexpressing 293T cells comparable with results of Ou et al. where ACE2 expression alone was also sufficient to induce cell-cell fusion without addition of exogenous protease (11), and also corroborated by a very recent report (41). Furthermore, we did not observe any effect on SARS2-S-mediated fusion activity upon inhibition of TMPRSS2 on target cells by the serine protease inhibitor Camostat when ACE2 was present. Together, these results imply that proteolytic activation by TMPRSS2 may not be a limiting factor for cell-cell fusion in 293T cells. A recent report demonstrated that upon co-transfection of spike, ACE2, and TMPRSS2, TMPRSS2 accelerates fusion. The size of the resulting syncytia only showed a TMPRSS2 dependency within the first 12h but was independent after 24h in that report (42), which is compatible with our observations of efficient cell-cell-fusion without TMPRSS2 in 293T. While SARS1-S-mediated cell-cell fusion was also weakly activated when ACE2 was expressed alone, activation was much higher in the presence of TMPRSS2, indicating stronger dependence of SARS1-S on TMPRSS2, compatible with the monobasic S1/S2 cleavage site in the SARS-CoV spike protein. Surprisingly, we even observed maximal activation with overexpression of only TMPRSS2, indicating that SARS1-S-mediated cell-cell fusion is mostly protease- and not ACE2-driven. In line with this observation, SARS1-S-mediated cell-cell fusion was clearly sensitive to Camostat, which reversed the TMPRSS2-mediated activation (Fig. 3 A).

Interestingly, mutational ablation of the S1/S2 cleavage site of SARS2-S rendered the mutated spike protein sensitive to inhibition by Camostat in the presence of ACE2 and TMPRSS2 (Fig. 3 A), suggesting that TMPRSS2 or a related protease is required for processing at the S2’ site to reach full activation when the S1/S2 site is not cleaved. In addition, in the absence of recombinantly expressed TMPRSS2, SARS2-S1/S2-mut was clearly impaired with regard to fusion activity (Fig. 2 A). Conversely, mutation of the S2’ priming site abrogated any effects of TMPRSS2 on SARS2-S-mediated fusion, e.g. when TMPRSS2 alone was provided by means of recombinant expression (Figs. 3 A, 6 A), or when TMPRSS2 was inhibited by Camostat (Fig. 6 E). It should be noted that the SARS2-S S2’ mutants were still fusogenic in the presence of high levels of ACE2 receptor (Figs. 2 A, 3 A, and 6 A, E), in the case of the GH and HH S2’ mutants even at moderate ACE2 levels, and in the absence of TMPRSS2 with similar activity as wildtype (Fig. 6 A). The S2’ GH mutant was also efficiently incorporated in (Fig. 7 D) and able to drive infection of pseudotyped lentiviral particles (Fig. 7 E). With wt SARS2-S or SARS2-S1/S2-mut, but not with the SARS2-S S2’ mutants, recombinant expression of TMPRSS2 led to low but detectable fusion activity (Figs. 2 A, 6 A). Collectively, these findings identify the S2’ site as the primary target of TMPRSS2 for fusion activation.

Another observation was that the S2’-AA mutant, as observed in Figs. 1 C and 6 C, exhibited drastically reduced surface expression. In fact, a similar incorporation defect has been described in the literature for SARS-CoV (43). Whatever the reason for this defect, we were able to overcome it completely by replacing the S2’ motif KR with the amino acids GH, which restored surface expression (Fig. 6 C), processing into S1 and S2 subunits (Fig. 6 B), and particle incorporation (Fig. 7 D). The reasons for this phenomenon are unclear. Charge reversal of S2’ from KR to EE was definitely detrimental to activity, indicating that solubility may not be the critical point. As histidine may carry a positive charge depending on the local environment, our findings might hint at a requirement for at least one positive charge at this position.

Our results clearly demonstrate that cleavage at the S1/S2 site alone is not sufficient for fusion activity in the presence of ACE2 and requires additional processing at S2’ or another site (26). This has been established for particle entry (24), but it was not entirely clear for cell-cell fusion, as the pre-cleaved spike was clearly fusogenic also in conditions without exogenous protease activity in several reports (11, 12, 26, 42), which could have been interpreted as a cell-cell fusion-ready state after S1/S2 cleavage. While our initial attempts to block the fusion activity of wt SARS2-S and the S1/S2 mutant in the presence of ACE2 receptor but without TMPRSS2 were relatively unsuccessful, treatment with the metalloprotease inhibitor Batimastat reduced fusion by both wt (Fig. 5 A) and the S1/S2 mutant (Fig. 5 C), as well as fusion by the S2’ mutants (Fig. 6 E). These findings indicate that metalloproteases can activate SARS2-S, and that this activation occurs at least in part independently of the S1/S2 site and of the S2’ site, as both mutants were still Batimastat-sensitive. On the other hand, the S1/S2 mutant was clearly less active in the absence of TMPRSS2, indicating that matrix metalloproteases activate more efficiently when the S1/S2 site is present. These findings are in line with a very recent report describing similar observations using different inhibitors (41).

As SARS2-S did not require TMPRSS2 on target cells for robust cell-cell fusion, our attempts to test the impact of Bromhexine as a specific inhibitor of TMPRSS2 on SARS2-S-mediated fusion activity were somewhat artificial. Nevertheless, SARS1-S-mediated fusion was clearly enhanced by TMPRSS2, as was fusion by SARS2-S1/S2-mut, and both were inhibited by Camostat but not by Bromhexine. Therefore, our finding that Bromhexine specifically enhanced fusion of 293T cells in the presence of SARS2-S, ACE2, and TMPRSS2 is something that we cannot explain easily.

According to our results the Bromhexine-mediated enhancement was specific for SARS2-S and was not seen with VSV-G as fusion effector (Fig. 3 C), nor did we observe significant effects with the SARS2-S mutants or SARS1-S (Fig. 3 A). We observed some inhibition of SARS1-S-mediated fusion in the presence of 50μM Ambroxol (Fig. 3 A) and also with SARS2-S with longer incubation times (Fig. 3 C), which may hint at some activity of this substance against TMPRSS2, which would fit with the observation of an atypical autoproteolytic fragment of TMPRSS2 in the presence of Ambroxol. The observation of the paradoxical effect of Bromhexine in the presence of TMPRSS2 suggests that Bromhexine somehow modulates proteolytic processing. It is at the moment not clear by what mechanism of action Bromhexine modulates TMPRSS2 activity, if it does so, and we therefore cannot exclude that processing of some substrates is actually enhanced or altered instead of inhibited, as reported for several substrates (15, 44). Recently, another study also reported lack of inhibitory activity of Bromhexine against TMPRSS2 (30). Activity of Bromhexine against TMPRSS2-mediated receptor shedding, which could also explain our observations, was not observed, in contrast to Camostat that increased ACE2 expression levels in the presence of TMPRSS2 (Figs. 3 B, 5 B). This may explain the slight increase, even if not always statistically significant, in fusion activity that we observed in some experiments with SARS2-S in the presence of Camostat when ACE2 and TMPRSS2 were co-expressed (Figs. 3 A, 5 A).

Compared to another study (45), our fusion assay yields slightly different results, with SARS2-S-mediated fusion appearing less dependent on activation by TMPRSS2. This could be due to differences in the protocol. The study by Yamamoto et al. allowed only for very short contact times of 4h and used non-adherent 293T FT cells, whereas we co-cultured the cells for a longer time, which allows for extended contact between cells and may enable the action of matrix metalloproteases. Our finding that TMPRSS2 is not required for fusion is in line with several reports making the same observation (11, 24, 26, 41, 42, 46). In general, we observed a higher fusion activity with our SARS2-S1/S2-mut spike mutant than was observed with furin cleavage site mutants in previous studies (12, 26), but we observed this only when TMPRSS2 was recombinantly overexpressed together with ACE2. When only ACE2 or only TMPRSS2 were recombinantly expressed, SARS2-S1/S2-mut fusion activity was strongly impaired (Fig. 2 A). It should be noted, that we left the loop intact and only replaced the basic residues with alanine in our mutant whereas other groups deleted the loop structure, which may result in a less flexible conformation. Nevertheless, our mutational approach for ablating the furin cleavage site clearly rendered the spike protein more dependent on additional serine protease activity by recombinantly expressed TMPRSS2. This proteolytic activity was directed towards the S2’ site, as SARS2-S1/S2-mut fusion activity was dependent on TMPRSS2 and was significantly inhibited by Camostat (Fig. 3 A) in the presence of TMPRSS2.

Taken together, our results actually reconcile several seemingly conflicting observations by other groups. The strong reduction in fusion activity by mutation of the S1/S2 site observed in one study in Vero (12) is reflected in our experimental conditions with only TMPRSS2 and endogenous levels of ACE2 expression, whereas our findings of more or less normal fusion activity under conditions of high-level ACE2 and TMPRSS2 expression are similar to the findings of another group with ACE2 overexpressing cells and addition of Trypsin or HAT (26).

Overall, we propose that the dependence on S1/S2 cleavage, activity of TMPRSS2 or a related protease and receptor expression are to a certain degree interdependent, where one factor can at least partially compensate for another, e.g. more extensive proteolytic activation at S2’ can render the spike more fusogenic even with lower receptor levels, which was particularly observed for SARS1-S, and to a lesser degree for SARS2-S (Fig. 2). Similarly, Batimastat-sensitive metalloproteases can activate SARS2-S for cell-cell fusion (Fig. 5 A). This is partially dependent on the S1/S2 site, as SARS2-S1/S2-mut was still impaired in the absence of TMPRSS2 but completely independent of the S2’ site, as demonstrated by full fusion activity of the SARS2-S2’-GH spike mutant on ACE2 expressing 293T cells (Fig. 6 A).

According to our results the requirements for cell-cell fusion and virus-cell fusion differ: Additional TMPRSS2 activity drastically enhanced pseudotype entry into transfected 293T (Fig. 7 A) but was not needed for cell-cell fusion with identically transfected 293T cells (Figs. 2 A, 3 A). In addition, the matrix metalloprotease inhibitor Batimastat did not affect particle entry in the presence of the TMPRSS2 inhibitor Camostat, indicating that matrix metalloproteases can activate cell-cell fusion but not particle-cell fusion (Fig. 7 C, E), at least not in our experimental system. Similar observations were previously made for SARS-CoV (34). The interpretation of these results is complicated by the ability of virus particles to enter cells through both direct membrane fusion or through an endocytotic pathway, and by different pre-priming states of viral spike proteins depending on proteolytic activity in the producer cell (47). As activation of the spike protein is expected to differ between organ systems depending on the presence of different proteolytic activities, these processes ultimately need to be studied in appropriate tissue systems or animal models. It is tempting to speculate that the relative to SARS-CoV more relaxed requirements for cell-cell fusion with regard to proteolytic activation contribute to the broad organ tropism and neuroinvasion by SARS-CoV-2, as well as the clinically observed formation of extended syncytia (13). Irrespective of the role of cell-cell fusion in COVID-19, in light of the observed paradoxical activation of cell-cell fusion by Bromhexine and its lack of inhibitory activity against entry of SARS2-S-pseudotyped lentiviruses on TMPRSS2 expressing cells, we would at the moment caution against clinical use of Bromhexine for treatment or prophylaxis of COVID-19, at least at higher concentrations that aim at the inhibition of TMPRSS2. A recent, small randomized trial showed promising results for Bromhexine at 3x 8 mg per day combined with hydroxychloroquine (48), which should result in Bromhexine plasma concentrations in the range of 0.1 μM (49). We are fairly confident to postulate that these favorable results are unlikely due to inhibition of TMPRSS2, although we can’t fully exclude extremely weak activity. This view is supported by a recent study that found no effect of Bromhexine on TMPRSS2 activity (30). More likely, favorable patient outcomes are attributable to the beneficial effects of Bromhexine or its main metabolite Ambroxol on lung function, general defense mechanisms against airway infections, and inflammatory response (16–19, 50). Another recent study by Olaleye et al (21) specifically analyzed the effects of Bromhexine and Ambroxol on the interaction of ACE2 with the SARS-CoV-2 spike receptor binding domain (RBD), and reported a very peculiar behavior of these substances which in part may explain the paradoxic results of our fusion assays and would support a beneficial effect of low-dose Bromhexine, which is converted to Ambroxol *in vivo*: While Ambroxol weakly inhibited the ACE2-RBD interaction up to 100μM concentration, Bromhexine exhibited a biphasic behavior and was weakly inhibitory below 10μM but increased ACE2-RBD binding at higher concentrations in that study. Both substances were reported to weakly inhibit SARS-CoV-2-mediated cytopathic effect in culture (22), and Ambroxol was also shown to moderately impact replication of SARS-CoV-2 in that report (22), albeit on Vero cells and not lung cells. Our results suggest that Ambroxol can weakly inhibit spike-driven entry of lentiviral pseudotypes into Calu-3 cells at high but potentially attainable concentrations (Fig. 8 B), and our experiments with authentic SARS-CoV-2 on Calu-3 (Fig. 8 C, D) demonstrated a trend towards inhibition of replication by both Ambroxol and Bromhexine, with Bromhexine possibly being slightly more potent, but also more toxic (Figs. 4 C, 8 E). Thus, the specificity of Bromhexine-mediated inhibition is questionable. In sum, it seems likely that Ambroxol acts weakly on TMPRSS2, which would explain its modest but significant effect on TMPRSS2-mediated activation of SARS1-S-mediated fusion (Fig 3 A). It should be noted that replication of the authentic virus can be influenced at numerous points, not necessarily only during entry, and that effects can be amplified over several replication cycles. Of course, compared to the potency of Camostat, the effect of both substances is marginal. Nevertheless, Ambroxol can be administered in high doses of 1g and more intravenously (19) or orally (51) and reportedly accumulates strongly in lung tissue (52). Thus, Ambroxol, which exhibited a trend towards inhibition of SARS2-S-mediated entry and fusion in several assays without enhancing effects as were observed with Bromhexine at high concentrations may represent an interesting option for supportive therapy at higher dosage, in particular as it is a proven therapeutic for antenatal respiratory distress syndrome (53) and has shown efficacy in the treatment of radiation-induced lung injury (50).

## MATERIAL AND METHODS

### Cell culture

All cell lines in this study were incubated at 37°C and 5% CO_2_. 293T (a kind gift from Vladan Rankovic and originally purchased from the ATCC, Göttingen) and Calu-3 cells (a kind gift from Stefan Ludwig) were cultured in Dulbecco’s Modified Eagle Medium (DMEM), high glucose, GlutaMAX, 25mM HEPES (Thermo Fisher Scientific) supplemented with 10% fetal calf serum (FCS) (Thermo Fisher Scientific), and 50μg/ml gentamycin (PAN Biotech). For Calu-3 cells 1mM Sodium-Pyruvate (Thermo Fisher Scientific) was added. For seeding and sub-culturing of cells the medium was removed, the cells were washed with PBS (PAN-Biotech) and detached with Trypsin (PAN-Biotech). All transfections were performed using PEI (Polysciences) in a 1:3 Ratio (μg DNA/μg PEI) mixed in OptiMEM. The cell viability assay with Calu-3 (Fig. 8 E) was performed as described previously (7); unlike for the other assays in this series of experiments Bromhexine and Ambroxol were used in the form of commercial cough suppressants (Bromo 12mg/ml, Krewel Meuselbach, and Mucosolvan 30mg/5ml, Sanofi-Aventis).

### Plasmids

Expression plasmids for pQCXIPBL-hTMPRSS2 (54), pCG1-SARS-2-S_humanized (55), pCG1-ACE2 (7) and pCG1-SARS S (56) are described elsewhere. For generation of pVAX1-SARS2-S the codon-optimized sequence encoding the spike protein of SARS-CoV-2 was amplified by PCR and cloned into the pVAX1 backbone. psPAX2 and pMD2.G were a gift from Didier Trono (Addgene plasmid # 12260, Addgene plasmid # 12259) and pLenti CMV GFP Neo (657-2) was a gift from Eric Campeau & Paul Kaufman (Addgene plasmid # 17447). Expression plasmids SARS2-S2’-AA, SARS2-S1/S2-mut, and SARS2-D614G were generated from pCG1_SL-Cov_Wuhan-S_humanized SARS2-S by PCR based mutation of the SARS2-S S1/S2 and the S2’ cleavage site using around-the-horn PCR mutagenesis using S7 Fusion PCR (Biozym) or Phusion PCR, T4 PNK and Quick ligase (all from New England Biolabs) and using the following primers: primers S1-S2 AAAA mut for V2 (CTGCCTCTGTGGCCAGCCAGAGCATC), S1-S2 AAAA mut rev V2 (CAGCGGCGGGGCTGTTTGTCTGTGTCTG), S2 to AA mut_Forward (GCCAGCTTCATCGAGGACCTGCTG) and S2 to AA mut_Reverse (AGCGCTGGGCTTGCTAGGATCGG), SARS2S R815 H for (CACAGCTTCATCGAGGACCTGCTG), SARS2S K814H rev (GTGGCTGGGCTTGCTAGGATCGG), SARS2S R815E for (GAGAGCTTCATCGAGGACCTGCTG), SARS2S K814E rev (CTCGCTGGGCTTGCTAGGATCGG), SARS2S R815E for (GAGAGCTTCATCGAGGACCTGCTG), SARS2S R815S for (AGCAGCTTCATCGAGGACCTGCTG), SARS2S K814G rev (TCCGCTGGGCTTGCTAGGATCGG), D614G for aroundthehorn (GCGTGAACTGTACCGAAGTGCC), D614G rev aroundthehorn (CCTGGTACAGCACTGCCACCTG). Sequence integrity was verified by sequencing of the coding region. Plasmid pCG1-SARS2-S_S2’mut contains a silent G to T mutation in the codon for leucine 441.

Expression plasmids pVAX1-SARS2-S_S2’-GH, pVAX1-SARS2-S1/S2-mut and pVAX1-SARS2-S_D614G were generated from pVAX1-SARS2-S by PCR-based mutation in a similar manner. The Gal4-Luc reporter plasmid encoding firefly luciferase under the control of an activator sequence that binds the Gal4 transcription factor has been described elsewhere (34). The Gal4 DNA binding domain VP16 fusion plasmid corresponds to Genbank identifier X85976. The TurboGFP-Luciferase fusion reporter gene was constructed using Gibson Assembly Master Mix (New England Biolabs) to insert the TurboGFP open reading frame with a Ser-Gly-Ser-Gly Linker in front of the Met codon of the luciferase open reading frame. Before assembly, the two fragments were generated using Phusion PCR (New England Biolabs) by amplifying the TurboGFP open reading frame from the vector pGIPZ (Thermo Scientific Open Biosystems) using the primers TurboGFP for Gal4Luc before ATG ov (GGTACTGTTGGTAAAATGGAGAGCGACGAGAGC) and TurboGFP rev (TTCTTCACCGGCATCTGCATC), and the Gal4-Luc backbone by amplification with primer Gal4Luc before ATG rev (TTTACCAACAGTACCGGAATGC) and primer Luc for SGSG TurboGFP overhang (GATGCAGATGCCGGTGAAGAAAGCGGTAGCGGTATGGAAGACGCCAAAAACATAAAG).

The pLentiCMV-TurboGFP::Luciferase fusion reporter gene was constructed using Gibson Assembly Master Mix (New England Biolabs) to exchange the insert in pLenti-CMV-Blast-EphA7-Strep (described elsewhere (57)) with the TurboGFP::Luc open reading frame without the Strep-Tag, the two fragments were generated using CloneAmp HiFi PCR Premix (Takara Bio) by amplifying the TurboGFP::Luc open reading frame from the Vector Gal4-TurboGFP-Luc using the primers GA_TurboGFP::Luc_pLentiBlast-StrepOneOv_For (acaaaaaagcaggctccaccATGGAGAGCGACGAGAGC) and GA_TurboGFP::Luc_pLentiBlast-StrepOneOv_Rev (tgtggatggctccaagcgctTTACAATTTGGACTTTCCGCC), and the pLenti-CMV-Blast-EphA7-Strep backbone by amplification with primer pLenti attB1 rev at ATG (CATGGTGGAGCCTGCTTTTTTGTAC) and OneStrep for (AGCGCTTGGAGCCATCCAC).

### Western blot

Protein expression was analyzed by polyacrylamide gel electrophoresis on 8%-16% precast gradient gels (Thermo) and Western blotting using antibodies to ACE2 (AF933, R&D Systems), c-Myc-epitope (clone 9E10, Santa Cruz Biotechnology), SARS spike (NB100-56578, Novus Biologicals), HIV-1 Gag p24 (clone 749140, R&D), and GAPDH (GenScript) in NETT-G (150 mM NaCl, 5mM EDTA, 50 mM Tris, 0.05% Triton X-100, 0.25% gelatin, pH 7.5) and donkey anti-mouse horseradish peroxidase (HRP)-coupled (Dianova), goat anti-rabbit HRP-coupled (Life Technologies) or rabbit anti-goat HRP-coupled (Proteintech) secondary antibody in 5% dry milk powder in PBS with 0.05% Tween 20. Imaging was performed using Immobilon Forte substrate (Merck) on an INTAS ECL ChemoCam system.

### Flow cytometry

293T cells were transfected with the respective spike expression constructs. On day two post transfection, the cells were harvested by gentle pipetting in PBS and were fixed in 2% methanol-free formaldehyde in PBS for 15 min. The cells were then washed once in PBS and then incubated in 10% FCS in PBS for 30 min to block non-specific binding. The cells were then incubated in either convalescent serum at 1:1000 dilution or soluble ACE2-Fc fusion protein at 2 ng/μl, both described elsewhere (58), for 1h in 10% FCS in PBS, followed by one wash in a large volume of PBS and then incubation with Alexa647-coupled anti-human secondary antibody (Thermo Fisher Scientific) at 1:200 in 10% FCS in PBS. The RRV gHΔ21-27-Fc fusion protein, which was used as a control protein, was generated from RRV 26-95 gH-Fc (59) by deletion of the codons for amino acid 21-27, which are important for receptor binding (28), and was produced analogous to the gH-Fc protein in Hahn et al. 2013 (59). The cells were then washed once in a large volume of PBS and post-fixated in 2% PFA in PBS before analysis on an LSRII flow cytometer (BD Biosciences). Data was analyzed using Flowing software (version 2.5) and GraphPad Prism, version 6, for Windows (GraphPad Software). COVID-19 convalescent serum was collected previously (58) in accordance with ethical requirements (ethics committee UK Erlangen, license number AZ. 174_20 B).

### Fusion assay

293T target cells were seeded in a 48-well plate at 50 000 cells/well and transfected with Vp16-Gal4 (Fig. 3 C) or Gal4-TurboGFP-Luciferase expression plasmid (Gal4-TurboGFP-Luc, all other experiments) as well as expression plasmids for ACE2 and TMPRSS2 as indicated. In case only ACE2 or TMPRSS2 were transfected the missing amount of DNA was replaced by empty vector. 293T effector cells were seeded in a 10 cm dish at 70-80% confluency and transfected with either the Vp16-Gal4 (all experiments except Fig. 3 C) or Gal4-Luciferase (Fig. 3 C) expression plasmid as well as expression plasmids for SARS2-S, SARS2-S1/S2-mut, SARS2-S2’-AA, SARS2-S2’-GH, SARS2-S2’-HH, SARS2-S2’-EE, SARS2-S2’-ES, SARS1-S, VSV-G glycoproteins or pcDNA6/V5-HisA (Thermo). For effector cell pre-incubation experiments, the medium of effector-cells was changed to Bromhexine hydrochloride (Merck), Ambroxol hydrochloride (Merck), Camostat mesylate (Tocris), Batimastat (Merck), AEBSF (Merck), EDTA (Merck), EGTA (Merck), 100x Cocktail Set V, Animal-Free - Calbiochem (Merck) or Decanoyl-RVKR-CMK (Merck) containing medium at final concentration 6h after transfection. 24h after transfection, target cells were pre-incubated with Bromhexine hydrochloride (Merck), Ambroxol hydrochloride (Merck), Camostat mesylate (Tocris), Batimastat (Merck), AEBSF (Merck), EDTA (Merck), EGTA (Merck) or Decanoyl-RVKR-CMK (Merck) for 30 min at the indicated concentration. Effector cells were then added to the target cells in a 1:1 ratio reaching the final inhibitor concentration. After 24-48h GFP-fluorescence was detected using a Vert.A1 Fluorescence Microscope and ZEN-Software (Zeiss), luciferase activity was analyzed using the PromoKine Firefly luciferase Kit or Beetle-Juice Luciferase Assay (PJKbiotech) according to manufacturer’s instructions and a Biotek Synergy 2 plate reader. Statistical analysis was performed using GraphPad Prism, version 6, for Windows (GraphPad Software).

### Production of lentiviral and pseudoparticles and pseudoparticle infection experiments

Lentiviral pseudoparticles were produced by transfecting 293T cells with expression plasmids for psPAX2, pLenti-CMV-GFP or pLentiCMV-TurboGFP::Luciferase and either SARS2-S variants (pVAX1-SARS2-S_S2’-GH, pVAX1-SARS2-S1/S2-mut and pVAX1-SARS2-S_D614G) or VSV-G (pMD2.G Addgene #12259). The cell culture supernatants were harvested 24-72 h post transfection followed by addition of fresh media and again after 48-72h. The supernatants were passed through a 0.45μm CA-Filter, and the SARS2-S pseudoparticles were concentrated via low speed centrifugation at 4°C for 16h at 4200xg. For detection of particle incorporation virus supernatant was further concentrated by centrifugation at 4°C for 2h at 21000xg on 5% Optiprep (Merck), the supernatant was removed and pellet was resuspended and subjected to Western Blot analysis. The SARS-CoV-2 spike and VSV-G lentiviral pseudoparticles were used to transduce 293T transfected with TMPRSS2 and ACE2 expression plasmids or Calu-3 cells. 48 h after transfection with control or ACE2 and TMPRSS2 expression plasmids, the pseudoparticles were added to the cells pre-incubated with inhibitors Bromhexine hydrochloride (Merck), Ambroxol hydrochloride (Merck), Camostat mesylate (Tocris), Batimastat (Merck), AEBSF (Merck) and E64-d (Biomol) for 30 min at twice the indicated concentration, the final concentration was reached after addition of the inoculum. Cells transduced with pLenti-CMV-GFP pseudoparticles were harvested 48 h after transduction using trypsin. Bald particles from 293T cells that were transfected with empty vector instead of glycoprotein expression plasmids and the lentiviral packaging system were used as background control for normalization. Trypsin activity was inhibited by adding 5% FCS in PBS, and after washing with PBS the cells were fixed with 4% formaldehyde (Roth) in PBS. The percentages of GFP-positive cells were determined using a LSRII flow cytometer, and at least 10000 cells were analyzed. Cells transduced with pLentiCMV-TurboGFP::Luciferase pseudoparticles were lysed after 48h with Luciferase-Lysis Buffer (Promega) and detected using Beetle-Juice Luciferase Assay according to manufacturer’s instructions and a Biotek Synergy 2 platereader. Statistical analysis was performed using GraphPad Prism 6.

### SARS-CoV-2 infections

Primary SARS-CoV-2 isolate ER-PR2 was a kind gift from Klaus Überla, Erlangen, and was originally isolated on Vero cells. The virus stock was then grown on Calu-3 cells in EMEM + 2% FCS + Penicillin/Streptomycin. Virus containing supernatant was harvested after CPE was clearly visible, and the supernatant was cleared by low-speed centrifugation at 1200 rpm for 10min before passage through a 0.2μm syringe filter (Mini-Sart, Sartorius). Virus stocks were aliquoted in 200μl aliquots and stored at −150°C. Infectivity was determined by the method of Reed and Muench (60) at 10^6.1^TCID50/ml. Calu-3 cells were seeded one day (first experiment) or two days (other two experiments) before infection and approx. 100000 Calu-3 cells were infected at a multiplicity of infection (MOI) of approximately 0.002 in a 96-well plate in triplicates. The cells were pre-incubated with the respective inhibitors in 50μl at twice the concentration for ~1.5h, the virus was then added in 50μl medium. Total RNA from the cells and the culture supernatant was harvested 20h (experiment 3) and 24h (experiments 1&2) post infection.

### RNA isolation, cDNA synthesis, and RT-qPCR

RNA was isolated using the Direct-zol RNA Miniprep Plus Kit (Zymo) according to the manufacturer’s instructions. For quantification of viral RNA in infected cultures, the cells and cellular supernatant in a volume of 100μl were lysed and inactivated by addition of 300μl TRI reagent (Zymo). RT-qPCR on viral genomes was performed using the N1 CDC primer set from IDT (2019-nCoV_N1-F GACCCCAAAATCAGCGAAAT and 2019-nCoV_N1-R TCTGGTTACTGCCAGTTGAATCTG, both 500nM, 2019-nCoV_N1-P FAM-ACC CCG CAT TAC GTT TGG TGG ACC-BHQ1, 125nM) and SensiFAST Probe Hi-ROX One-Step Kit (Bioline) according to the manufacturer’s instructions in a 20μl reaction with 5μl sample. All RT-qPCR reactions were performed in technical duplicates on a StepOne Plus (Thermo) realtime cycler. PCR conditions were 45°C for 10min, 95°C for 2min, and then 45 cycles 95°C for 5sec followed by 55°C for 20sec. To determine the PCR efficiency across the whole dynamic range, a 7-step 10-fold dilution series with the H_2_O-treated SARS-CoV-2-infected Calu-3 sample was performed. These datapoints with the undiluted sample set to 1 was approximated by an exponential function using Microsoft Excel 2020. The measured PCR efficiency was additionally fitted by multiplication with a constant factor to match our RNA standard (Charite, Berlin), which was only available at 50, 500, and 5000 copies, which confirmed our approach, but was not used for relative quantification. Fit was performed by minimizing the sum of the squared relative deviations from the standard concentrations with an exactness of two digits.

For quantification of cellular TMPRSS2 and GAPDH expression cDNA synthesis and qPCR were performed according to the manufacturer’s instructions using the SensiFast cDNA kit and SensiFAST SYBR qPCR kit (both from Bioline). The qPCR was run on a StepOnePlus realtime PCR cycler (Thermo) and analyzed using the StepOne Software, which was also used to calculate ΔΔCt values and error estimates for TMPRSS2 expression. TMPRSS2 mRNA was detected using primer set Hs.PT.58.39408998 (IDT) (forward primer GTCAAGGACGAAGACCATGT, reverse primer TGCCAAAGCTTACAGACCAG). GAPDH mRNA was detected using primers GAPDH_Hs-Mm_s (CTTTGGTATCGTGGAAGGACTC) and GAPDH_Hs-Mm_as (GTAGAGGCAGGGATGATGTTC). Amplifications with Ct above 35 and non-matching melting curve were scored as not detected.

## ACKNOWLEDGEMENTS

We thank Stefan Pöhlmann and Markus Hofmann for sharing reagents and for critical reading of the manuscript and helpful discussions. We thank Klaus Überla for sharing SARS-CoV-2 ER-PR2. We also thank Armin Ensser and Florian Full for helpful discussions.

